# Multiple links between 5-methylcytosine content of mRNA and translation

**DOI:** 10.1101/2020.02.04.933499

**Authors:** Ulrike Schumann, He-Na Zhang, Tennille Sibbritt, Anyu Pan, Attila Horvath, Simon Gross, Susan J Clark, Li Yang, Thomas Preiss

## Abstract

5-methylcytosine (m^5^C) is a prevalent base modification in tRNA and rRNA but it also occurs more broadly in the transcriptome, including in mRNA, where it serves incompletely understood molecular functions. In pursuit of potential links of m^5^C with mRNA translation, we performed polysome profiling of human HeLa cell lysates and subjected RNA from resultant fractions to efficient bisulfite conversion followed by RNA sequencing (bsRNA-seq). Bioinformatic filters for rigorous site calling were devised to reduce technical noise. We obtained ∼1,000 candidate m^5^C sites in the wider transcriptome, most of which were found in mRNA. Multiple novel sites were validated by amplicon-specific bsRNA-seq in independent samples of either human HeLa, LNCaP and PrEC cells. Furthermore, RNAi-mediated depletion of either the NSUN2 or TRDMT1 m^5^C:RNA methyltransferases showed a clear dependence on NSUN2 for the majority of tested sites in both mRNAs and noncoding RNAs. Candidate m^5^C sites in mRNAs are enriched in 5’UTRs and near start codons, and are commonly embedded in a local context reminiscent of the NSUN2-dependent m^5^C sites found in the variable loop of tRNA. Analysing mRNA sites across the polysome profile revealed that modification levels, at bulk and for many individual sites, were inversely correlated with ribosome association. Altogether, these findings emphasise the major role of NSUN2 in making this mark transcriptome-wide and further substantiate a functional interdependence of cytosine methylation level with mRNA translation.

## Introduction

Cells across all domains of life have an impressive ability to ‘decorate’ their RNAs post-transcriptionally; the MODOMICS database (Boccaletto et al. 2018) currently lists over 170 known types of chemically modified ribonucleosides and over 360 different proteins involved in RNA modification. This chemical diversity abounds among the noncoding (nc)RNAs involved in translation, and the transfer (t)RNA research field in particular has been a principal source of RNA modification discovery for decades (Helm and Alfonzo 2014). By contrast, despite early indications, *e.g.* (Desrosiers et al. 1974; Perry and Kelley 1974; Dubin and Taylor 1975), technological barriers hindered research into the presence and specific distribution of modified nucleosides within messenger (m)RNAs and other ncRNAs. This changed when next generation sequencing methods were adapted, first to detect RNA editing events, reviewed in (Ramaswami and Li 2016), and soon after to map 5-methylcytosine (m^5^C) and *N*6-methyladenosine (m^6^A) in a transcriptome-wide fashion (Dominissini et al. 2012; Meyer et al. 2012; Squires et al. 2012).

Expansion in scope and refinement of such methods (Schaefer et al. 2017; Linder and Jaffrey 2019; Motorin and Helm 2019) have now produced maps of several modifications in tissues and cells of diverse origins (Fray and Simpson 2015; Burgess et al. 2016; Marbaniang and Vogel 2016; Kennedy et al. 2017; Shen et al. 2019), spawning the term ‘epitranscriptomics’ to mainly (but not exclusively) refer to research into the function of modified nucleosides in mRNA (Saletore et al. 2012). Challenges exist not only in the accurate detection of these generally sparse modifications but also in ascribing molecular, cellular and organismic functions to these ‘epitranscriptomic marks’—in mRNA (Sibbritt et al. 2013; Peer et al. 2017; Nachtergaele and He 2018) as well as in ncRNA (Shafik et al. 2016; Jacob et al. 2017). Here, in analogy to DNA epigenetics, the concept of RNA modification ‘writers, readers, and erasers’ has become an influential, if not always perfectly suited, guide to thinking in the field (Liu and Pan 2015; Roundtree and He 2016; Meyer and Jaffrey 2017; Shi et al. 2019; Zaccara et al. 2019).

How this plays out can be seen with the long suspected (Sommer et al. 1978) and recently substantiated mRNA destabilising effects of m^6^A. The modification is added co-transcriptionally in the nucleus by the m^6^A ‘writer’ complex, which includes the methyltransferase (MTase) METTL3 (methyltransferase-like protein 3) (Liu et al. 2014), while it can also be ‘erased’ again by the demethylase ALKBH5 (AlkB homolog 5) (Zheng et al. 2013). Several proteins have been shown to bind or ‘read’ m^6^A, including members of the YTH domain-containing family (YTHDF). Among them, YTHDF2 is known to promote mRNA decay in the cytoplasm (Wang et al. 2014; Du et al. 2016). However, in addition to its role in turnover, m^6^A has also been implicated in mRNA processing, export and translation, reviewed in (Zaccara et al. 2019), as well as in editing (Xiang et al. 2018). Thus, m^6^A illustrates what can be expected of RNA modifications more broadly, namely that they might have diverse, context-dependent functions. Context relates to both, where the modification is found within an RNA (*e.g.* sequence, structure and modification level; proximity or overlap with other functional/regulatory RNA features) but also the broader cellular milieu (*e.g.* availability of ‘readers’ and their downstream effectors) (Sibbritt et al. 2013; Shi et al. 2019).

m^5^C is present in multiple tRNAs, where it can influence the accuracy of translation and tRNA stability (Phizicky and Alfonzo 2010), thereby also affecting the formation of tRNA-derived small regulatory ncRNAs (Schaefer et al. 2010; Anderson and Ivanov 2014; Blanco et al. 2014). m^5^C sites are also found in ribosomal (r)RNA and they can affect ribosome biogenesis, stability and translational performance (Sloan et al. 2017). The eukaryotic m^5^C ‘writers’ are the seven members of the NOL1/NOP2/SUN domain (NSUN) MTase family and TRDMT1 (tRNA aspartic acid MTase 1; *a.k.a.* DNA MTase homologue 2, DNMT2). They have mostly been characterised as either targeting tRNA (NSUN2, 3 and 6; TRDMT1) or rRNA (NSUN1, 4 and 5) (Sibbritt et al. 2013; Bohnsack et al. 2019). Despite their seemingly ‘housekeeping’ functions, these MTases display complex expression patterns during development and disease, especially in cancer, and mutations in several of them cause human genetic disease (Chi and Delgado-Olguin 2013; Blanco and Frye 2014; Begik et al. 2019; Bohnsack et al. 2019). One explanation for their complex biology might be that these MTases modify additional substrates outside of the tRNA and rRNA realm. Indeed, NSUN7 was recently identified as a modifier of enhancer RNAs (Aguilo et al. 2016), but NSUN2 has also repeatedly been found to methylate sites outside of its purview (Squires et al. 2012; Hussain et al. 2013b; Khoddami and Cairns 2013; Yang et al. 2017; Chen et al. 2019; Huang et al. 2019; Sun et al. 2019). Two ‘readers’ of m^5^C have been reported, the mRNA export adapter ALYREF (Aly/REF export factor) (Yang et al. 2017) and the DNA/RNA-binding protein YBX1 (Y-box binding protein 1) (Chen et al. 2019; Yang et al. 2019b), implying certain molecular functions (see below). Finally, a potential route to ‘erase’ m^5^C from RNA is indicated by the presence of 5-hydroxymethylcytosine (hm^5^C), 5-formylcytosine and 5-carboxylcytosine in RNA, which represent intermediates in an oxidative demethylation pathway initiated by Ten-eleven translocation (TET) dioxygenases (Delatte et al. 2016; Huang et al. 2016; Miao et al. 2016; Zhang et al. 2016).

Transcriptome-wide m^5^C maps at variable depth are by now available for several tissues and cell lines of human/mouse (Squires et al. 2012; Hussain et al. 2013b; Khoddami and Cairns 2013; Blanco et al. 2014; Blanco et al. 2016; Amort et al. 2017; Legrand et al. 2017; Yang et al. 2017; Wei et al. 2018; Chen et al. 2019; Huang et al. 2019; Sun et al. 2019), zebrafish (Yang et al. 2019b), plant (Cui et al. 2017; David et al. 2017; Yang et al. 2019a), archaeal (Edelheit et al. 2013) and even viral (Courtney et al. 2019a; Courtney et al. 2019b) origin, persistently identifying sites with biased distribution in mRNAs and/or ncRNAs, reviewed in (Trixl and Lusser 2019). Several studies have further suggested regulatory roles for m^5^C in mRNAs. For example, it was shown in the context of leukemia that m^5^C in nascent RNA mediates formation of specific active chromatin structures (Cheng et al. 2018). m^5^C can also guide systemic mRNA transport in plants (Yang et al. 2019a), promote nuclear export of mammalian mRNA in conjunction with ALYREF (Yang et al. 2017) and enhance mammalian and zebrafish mRNA stability facilitated by YBX1 (Chen et al. 2019; Yang et al. 2019b). Further, a negative correlation was noted between translation of mammalian mRNAs and the presence of m^5^C sites transcriptome-wide (Huang et al. 2019). Finally, there is a body of work on individual mRNAs and their regulation by m^5^C at the levels of stability and translation in the context of cell proliferation and senescence (Wang 2016; Casella et al. 2019).

Different approaches based on high-throughput RNA-seq as a readout have been developed to map m^5^C. One is to perform an immunoprecipitation of cellular RNA fragments with anti-m^5^C antibodies (m^5^C-RIP) (Edelheit et al. 2013; Cui et al. 2017). Other approaches use enzyme-trapping, either by over-expressing a mutant MTase that cannot resolve the covalent enzyme-RNA intermediate (methylation iCLIP or miCLIP) (Hussain et al. 2013b), or by prior incorporation of 5-azacytidine into cellular RNA, which then covalently traps endogenous MTases to their substrates (Aza-IP) (Khoddami and Cairns 2013). Further, in analogy to epigenetic detection in DNA, resistance of m^5^C to conversion into uridine by bisulfite treatment of RNA has also been used (bisulfite RNA-seq, bsRNA-seq) (Squires et al. 2012; Amort et al. 2017; David et al. 2017; Legrand et al. 2017; Yang et al. 2017; Huang et al. 2019; Yang et al. 2019b). Imperfections of each method have been noted, for example, in m^5^C-RIP antibody specificity is crucial, while for miCLIP and Aza-IP sensitivity might not reach lower abundance targets. bsRNA-seq is not completely specific to m^5^C (*e.g.* it also detects hm^5^C) and is affected by incomplete conversion of unmodified cytosines due to RNA structure and the variability in reaction conditions. Stringency criteria to balance false negative against false positive site calls have also been set differently between studies, leading to, for example, drastically different estimates for sites in mRNA from a handful to thousands (Legrand et al. 2017; Linder and Jaffrey 2019). Regardless of chosen method, a redeeming feature might be that with greater insight into experimental limitations and with refinement of bioinformatic methods, new, better quality m^5^C epitranscriptomic maps can supersede early, pioneering attempts.

Here, we pursued the molecular roles of m^5^C in human mRNA with an emphasis on any link to translation. RNA isolated from multiple polysome profiling fractions was subjected to efficient bisulfite conversion followed by RNA sequencing (bsRNA-seq), rigorous candidate m^5^C site calling and validation. Bioinformatic analyses identified the sequence and structural context of sites, their preferred location along mRNA, and multiple correlative links to translation.

## Results

### Transcriptome-wide bsRNA-seq after separation by translation state

For polysome profiling, rapidly growing HeLa human cervical cancer cells (in biological triplicates B, C and E) were lysed in the presence of cycloheximide, lysates separated by ultracentrifugation through linear sucrose density gradients and multiple fractions taken (Clancy et al. 2007) (Figure 1A and Table S1 for parameters of each lysate). To monitor efficacy of separation on the gradients and reproducibility across replicates, the absorbance profile at 254 nm was recorded and the distribution of Ribosomal Protein L26 (RPL26) measured (Figure 1B,C). RNA was isolated from fractions, which were spiked with Renilla Luciferase (*R-Luc*) RNA transcribed *in vitro* (sequences shown in Table S2), and its integrity checked (Figures 1D, S1A). RNA fractions were then used in RT-qPCR (primers listed in Table S3) to establish the sedimentation behaviour of multiple cellular mRNAs (Figures 1E, S1B). To best capture the varying mRNA profiles and, therefore, different translation states, we pooled RNA samples into four final fractions for bsRNA-seq. Fraction 1 encompassed the small ribosomal subunit peak, fraction 2 included the large ribosomal subunit and monosomal peaks, whereas fractions 3 and 4 covered light and heavy polysomal peak regions, respectively (indicated by the blue boxes in Figure 1). Pooled bsRNA-seq fractions were prepared from each biological triplicate (Figure S1C), laced with the ERCC (External RNA Controls Consortium) spike-in mix (Baker et al. 2005), depleted of rRNA and subjected to bisulfite conversion (see Figure S1D for RNA integrity analyses before and after these steps). Libraries (termed **LibB1-4**, **LibC1-4** and **LibE1-4**) were prepared and subjected to Illumina HiSeq sequencing (bsRNA-seq).

**Figure 1:**
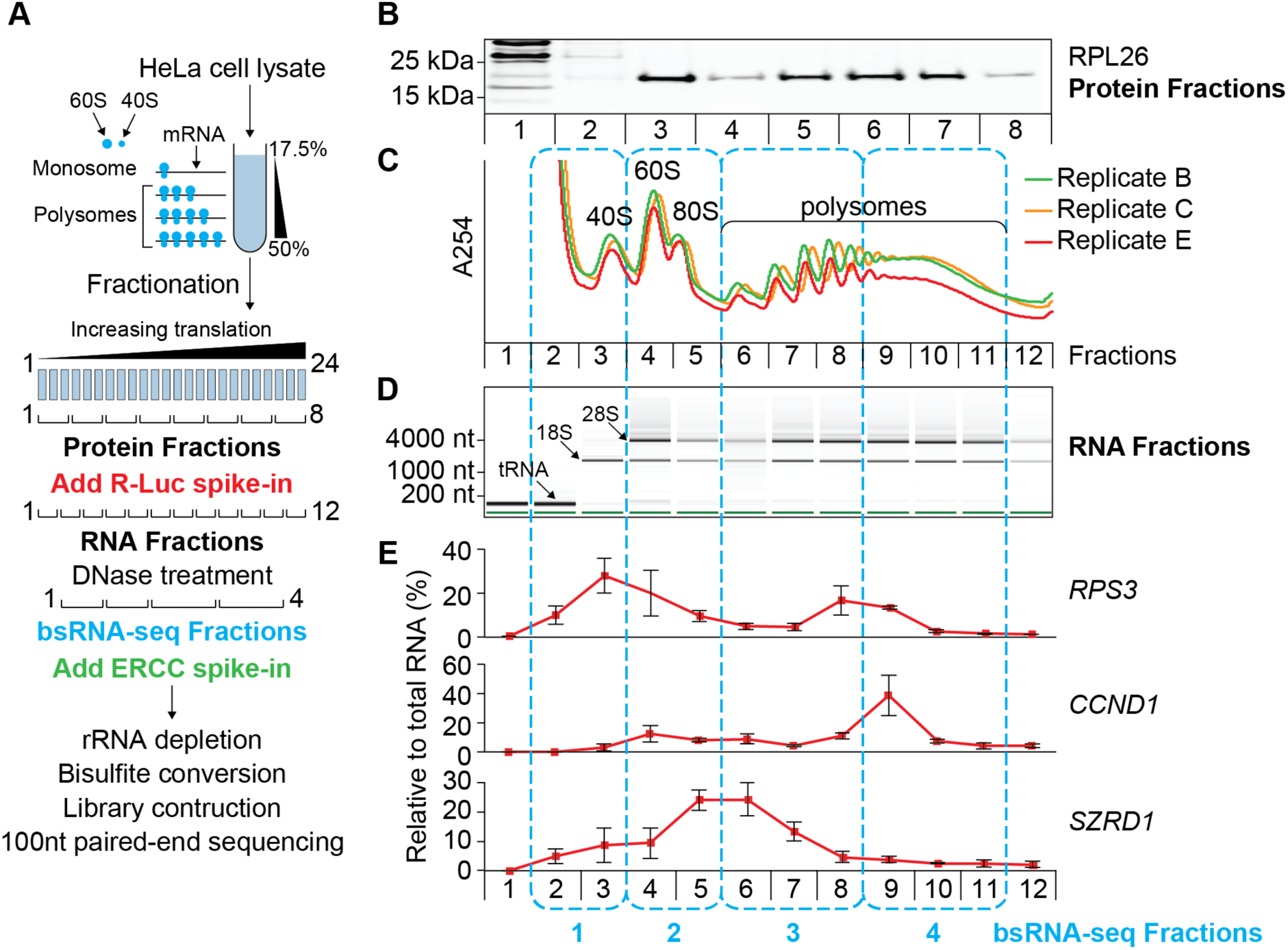
Workflow of polysome profiling and sample selection for bisulfite (bs)RNA-seq. HeLa cell lysates (three biological replicates, B, C and E) were separated by ultracentrifugation through linear sucrose density gradients. Twenty-four fractions per gradient were initially taken and further combined for subsequent analyses as indicated. Of particular note, the four pooled fractions chosen for bsRNA-seq are indicated by blue boxes. A: Principle of using sucrose density gradient ultracentrifugation to separate cellular mRNAs by increasing ribosome association (translation state; top) and scheme for merging the initial twenty-four fractions into different pools for downstream analyses including bsRNA-seq (bottom). First, subsamples were taken and three adjacent fractions each were merged to generate eight samples for Western blotting (**Protein Fractions**). Second, fractions were spiked with a Renilla luciferase (*R-Luc*) *in vitro* transcript and combined pairwise to generate twelve merged fractions (**RNA Fractions**). Total RNA was isolated, treated with DNase, assessed for RNA integrity and used for mRNA-specific RT-qPCR. Third, pools (**bsRNA-seq Fractions**) were made from selected RNA samples as follows: (pool 1: RNA fractions 2-3; pool 2: 4-5; pool 3: 6-8; pool 4: 9-11). 10μg total RNA from each of the four pools was spiked with the ERCC *in vitro* transcripts, depleted of rRNA, treated with sodium bisulfite prior to library construction and high-throughput Illumina sequencing. B: Distribution of Ribosomal Protein L26 (RPL26) across gradients. Protein fractions were subjected to western blotting against RPL26 (equal proportions of combined fractions were loaded). Blot of replicate B is shown as an exemplar of all biological replicates. C: Absorbance traces (254nm) across the three independent biological replicate gradients processed for bsRNA-seq. Total RNA was isolated from the twelve RNA fractions for downstream analyses. D: Distribution of tRNA and rRNA across gradients. RNA fractions were analysed by microfluidic electrophoresis (equal proportions of recovered RNA were loaded). The pseudo-gel image for replicate E is shown (see Figure S1A for data from all replicates). E: Distribution of representative mRNAs across gradients. mRNA levels in each RNA fraction were determined by RT-qPCR. Results for three mRNAs of different coding region length are shown: *RPS3* (ribosomal protein S3), *CCND1* (cyclin D1) and *SZRD1* (SUZ RNA binding domain containing 1) (see Figure S1B for further examples). mRNA levels per fraction were normalised to the level of a spike-in control, rescaled as percentage of total signal across all fractions, and are shown as mean across the three biological replicates with error bars indicating ± standard deviation.

### Transcriptome-wide identification of candidate m^5^C sites

bsRNA-seq yielded on average ∼55 million (M) read pairs per library after initial processing. Reads were mapped to ‘bisulfite-converted’ references as shown in Figure S2 and detailed in Methods. Of further note, the human reference genome was combined with the spike-in sequences, whereas dedicated references were used for rRNA and tRNA mapping. On average, ∼58% of reads were uniquely mapped to the genome (an additional ∼1.4%, ∼0.07%, ∼4.9% and ∼0.04% of reads mapped to the ERCC and *R-Luc* spike-ins, rRNA and tRNAs, respectively; mapping statistics are given in Table S4).

Next, we merged the four bsRNA-seq fraction libraries into one composite library per biological replicate (termed **cLibB**, **cLibC** and **cLibE**, *i.e.* N=3) to approximate a total cellular RNA analysis (Figure S2). Using the initial read mapping, we then assessed overall cytosine conversion for different RNA types. We saw near complete conversion (>99.8%) of both ERCC and *R-Luc* spike-ins, which are devoid of modified nucleosides and thus attest to the efficiency of the bisulfite conversion reaction (Figure 2A). C-to-T conversion was also very high across the transcriptome (∼99.7% for all annotated RNAs; ∼99.8% for transcripts of protein-coding genes), consistent with a rare occurrence of m^5^C sites in mRNA (Legrand et al. 2017; Yang et al. 2017; Chen et al. 2019; Huang et al. 2019; Sun et al. 2019). Conversion levels for tRNAs were lower (∼94.2%), as expected from the common presence of m^5^C sites in tRNAs. The conversion of rRNAs was intermediate (∼97.8%), which is inconsistent with the sparseness of m^5^C sites within mature rRNAs (only two sites known in 28S rRNA). We further visually inspected non-conversion at individual cytosine positions for multiple spike-in RNAs and tRNAs as well as for 28S, 18S and 5.8 rRNAs (Figure S3). Initial read mapping already reported high levels of non-conversion at known m^5^C positions within tRNAs as well as near complete conversion at other tRNA positions and at all cytosines within the spike-in transcripts (top panels in Figure S3A,B). By contrast, mapping to rRNAs suffered from incomplete cytosine conversion in multiple clusters (top panels in Figure S3C), particularly in 28S rRNA. This clustered non-conversion likely reflects the known sensitivity of the bisulfite reaction to secondary structure (Goddard and Schulman 1972; Goodchild et al. 1975; Goddard and Maden 1976) and indicated additional filtering as necessary for high-confidence site calls.

**Figure 2:**
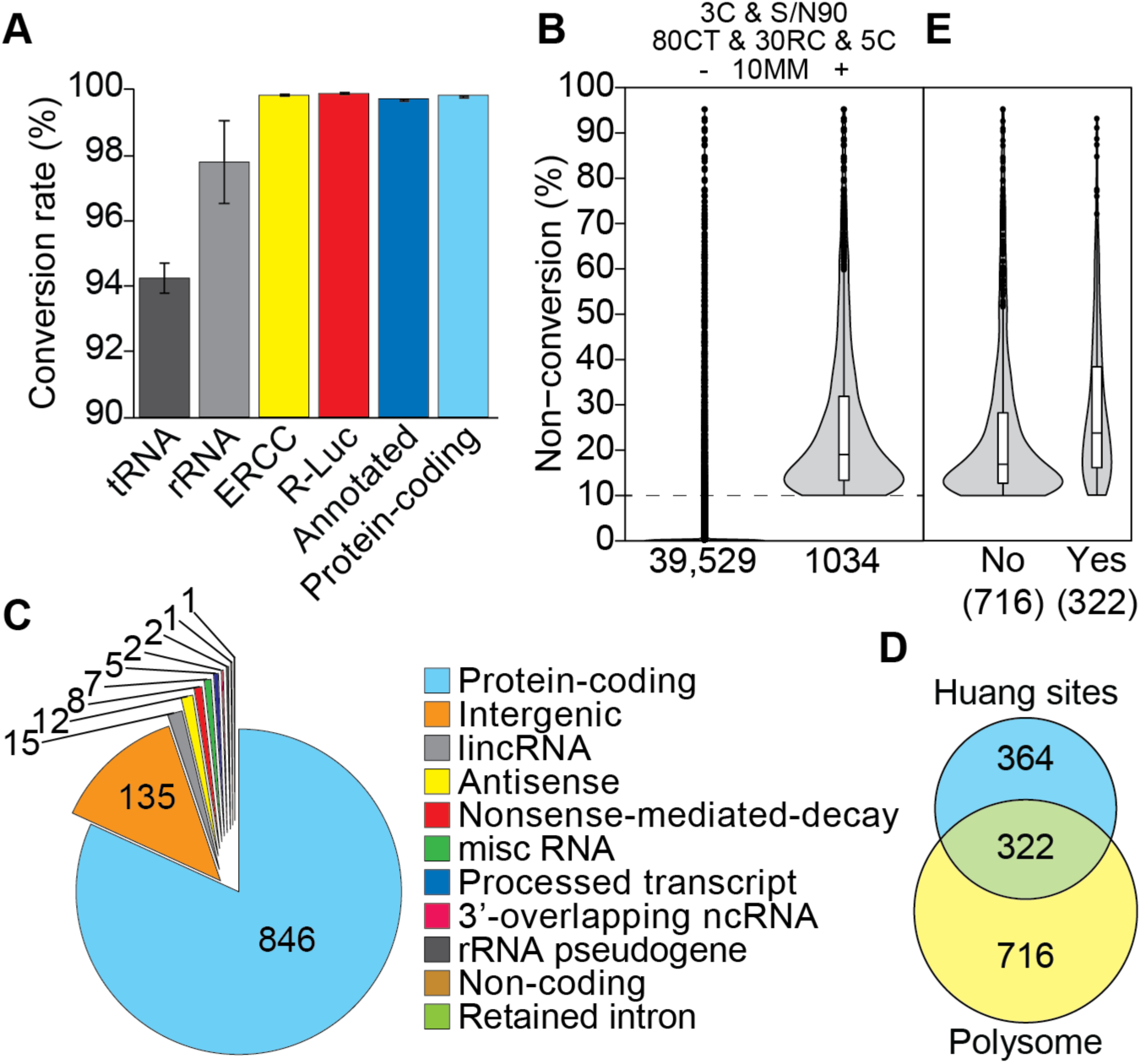
bsRNA-seq performance and transcriptome-wide candidate m^5^C site distribution. bsRNA-seq fraction library mapping data were combined into one composite dataset for each biological replicate (e.g. LibB1-4 yielded cLibB, see Figure S2). A: Cytosine conversion rate within different RNA types based on initial read mapping. tRNA and rRNA data are based on dedicated mapping to specialised references, others (ERCC and *R-Luc* spike-ins, all annotated transcripts and protein-coding transcripts [based on GENCODE v28]) are from mapping to the general reference genome (see Materials and Methods for details). Shown are mean values across the three biological replicates with error bars indicating ± standard deviation. B: Violin plots showing cytosine non-conversion range of transcriptome-wide m^5^C candidate sites, present in all three replicates and after 3C & S/N90, 80CT, 30RC and 5C filtering, either before (left) or after application of a ≥10% average non-conversion cut-off (10MM, right; 1,034 sites pass all filters, see main text for details). C: Distribution of 1,034 transcriptome-wide m^5^C candidate sites across RNA biotypes. Site annotation was according to the longest transcript variant recorded in GENCODE v28. Note that 84 of the 135 ‘intergenic’ sites are in fact annotated as tRNAs in RefSeq. D: Venn diagram showing overlap between 1,034 m^5^C candidate sites identified here and a set of 686 sites reported for poly-A-enriched HeLa cell RNA elsewhere (Huang et al. 2019). Huang et al. applied bsRNA-seq to HeLa control, NSUN2 knock-out and NSUN2-rescue samples. A union of called sites with at least 10% non-conversion in any of their three samples were used for this analysis. E: Violin plots showing cytosine non-conversion range of m^5^C candidate sites reported here after subdivision into those that do (Yes) or do not (No) overlap with the set reported by (Huang et al. 2019).

As a first step towards that, we removed reads that contain more than three non-converted cytosines from the initial read mapping (the **‘3C’** filter) (Edelheit et al. 2013; Blanco et al. 2014; Huang et al. 2019). While this did not affect the analysis of spike-in RNAs and most tRNAs (middle panels in Figure S3A,B), it noticeably reduced, but did not completely eliminate, clustered non-conversion for rRNAs (middle panels in Figure S3C). Thus, as also suggested by others (Huang et al. 2019), we further implemented a ‘signal-to-noise’ filter, that suppresses site calls at positions where less than 90% of mapped reads passed the 3C filter (**‘S/N90’**). Combined application of the 3C and S/N90 filters (3C & S/N90) did not change site calls for most tRNAs, although predictably, those with more than three genuine m^5^C sites were affected (bottom panels in Figure S3B). While detection of the two known m^5^C sites at position 3,761 and 4,417 in 28S rRNA benefited from the 3C filter, they were both flagged as unreliable by the S/N90 filter (compare middle and bottom panels in Figure S3C, as well as the ‘zoomed plots’, Table S5A). On balance, we accepted this tendency for false negative site calls, at least in tRNA and rRNA, in favour of suppressing false positive calls transcriptome-wide.

Additional criteria we implemented for inclusion as a candidate m^5^C site: coverage above thirty reads (**‘30RC’**), at least 80% of bases identified are cytosine or thymidine (**‘80CT’**) and a minimal depth of at least five cytosines (**‘5C’**). Reproducibility of sites called in this way was high across the biological triplicates (R^2^ >0.95; Figure S4A). Nevertheless, most called sites had very low cytosine non-conversion (e.g. ≤1% on average for 35,090 of a total 39,529; Figure 2B). Thus, to select sites likely to be ‘biologically meaningful’, we finally required an average non-conversion level across replicates of at least 10% (**‘10MM’**). This identified 1,034 high-confidence candidate m^5^C sites in the transcriptome-wide mapping (Figure 2B, Table S5B), with only a minority of sites showing very high levels of non-conversion (e.g. ∼5% had levels above 60%). 322 of these 1,034 sites were also included in a set of HeLa cell m^5^C sites reported recently based on bsRNA-seq with similar mapping and site selection strategies (Huang et al. 2019) (Figure 2D). The underlying concordance of the datasets is likely higher, as our data has around four-fold greater depth of coverage and overlapping the two site lists further suffers from non-conversion thresholding effects (leading to an exaggerated lack of overlap between sets around the 10% cut-off; Figure 2E). Given their critical impact on data clean-up, the specific effects of the 3C and S/N90 filters on numbers of called sites in different RNA types is shown in Figure S4B (with all other filters left in place). Notably, the great majority of candidate m^5^C sites (846 of 1,034) was found in transcripts of protein coding genes (Figure 2C); these became the main focus of further analyses, as detailed below.

### Candidate m^5^C sites in tRNAs and other ncRNAs

Given that polysomal RNA was our source material, coverage for most ncRNAs was not expected to be high. Nevertheless, ∼18% of sites that passed all our criteria mapped to ncRNA biotypes (Figure 2C). Notable numbers were found in long intergenic ncRNAs (15 sites) and antisense transcripts (12 sites). These include sites in ribonuclease P RNA component H1 (*RPPH1*; *chr14:20,343,234*), small Cajal body-specific RNA 2 (*SCARNA2*; *chr1:109,100,508*), RNA component of signal recognition particle 7SL1 (*RN7SL1*; *chr14:49,586,869*) and two RN7SL pseudogenes, *RN7SL395P* (*chr8:144,785,379*) and *RN7SL87P* (*chr5:144,140,971*), as well as sites in the two vault RNAs, *vtRNAs1-1* (*chr5:140,711,344* and *chr5:140,711,359*) and *vtRNA1-2* (*chr5:140,718,999*), all of which have been previously reported (Squires et al. 2012; Hussain et al. 2013a; Khoddami and Cairns 2013). Additionally, we identified candidate sites that had not been specifically noted by previous studies, in NSUN5 pseudogene 2 (*NSUN5P2; chr7:72,948,484*), telomerase RNA component (*TERC; chr3:169,764,738*) and nuclear paraspeckle assembly transcript 1 (*NEAT1*; *chr11:65,425,307*). Although it did not fulfil our coverage criteria, we also saw evidence of non-conversion in *SNORD62B* (*chr9:131,490,541*). The sites in *RPPH1*, *SCARNA2*, *NSNU5P2* and *SNORD62B* were further validated in independent biological samples (see below).

135 sites were located in intergenic regions, according to GENCODE v28 annotation, however, 84 of these reside within tRNAs according to RefSeq annotation. We systematically identified sites in tRNAs from reads that uniquely mapped within the processed tRNA sequence coordinates according to our bespoke pre-tRNA reference (see Methods; ∼175,000 reads per cLib/replicate). tRNA coverage in our data is comparatively low as the bulk of tRNAs sediment near the top of the gradient (Figure 1C), a region we did not include in our bsRNA-seq fraction selection. Nevertheless, we identified a total of 119 candidate m^5^C sites in 19 tRNA iso-decoders (Table S5C). These sites typically show high cytosine non-conversion level (see examples in Figure S3C) and they are near exclusively located in the anticipated tRNA secondary structure positions (Figure S4C). The abundant identification of the major known tRNA sites confirms the reliability of our dataset.

### Validation of candidate m^5^C sites and their NSUN2-dependence

We employed a targeted approach termed amplicon-bsRNA-seq to confirm the presence of selected sites in independent biological samples. It uses RT-PCR to amplify specific transcript regions after bisulfite conversion and purified amplicons are sequenced using Illumina MiSeq technology. Altogether, we report amplicon-bsRNA-seq for twenty-six different RNAs in this study, including two *R-Luc* spike-ins and two tRNAs (as controls; Figure S5) as well as seventeen mRNAs (two sites in the 5’UTR, nine in the CDS and six in the 3’UTR) and five ncRNAs (Figures 3,S6; see also Table S6).

**Figure 3:**
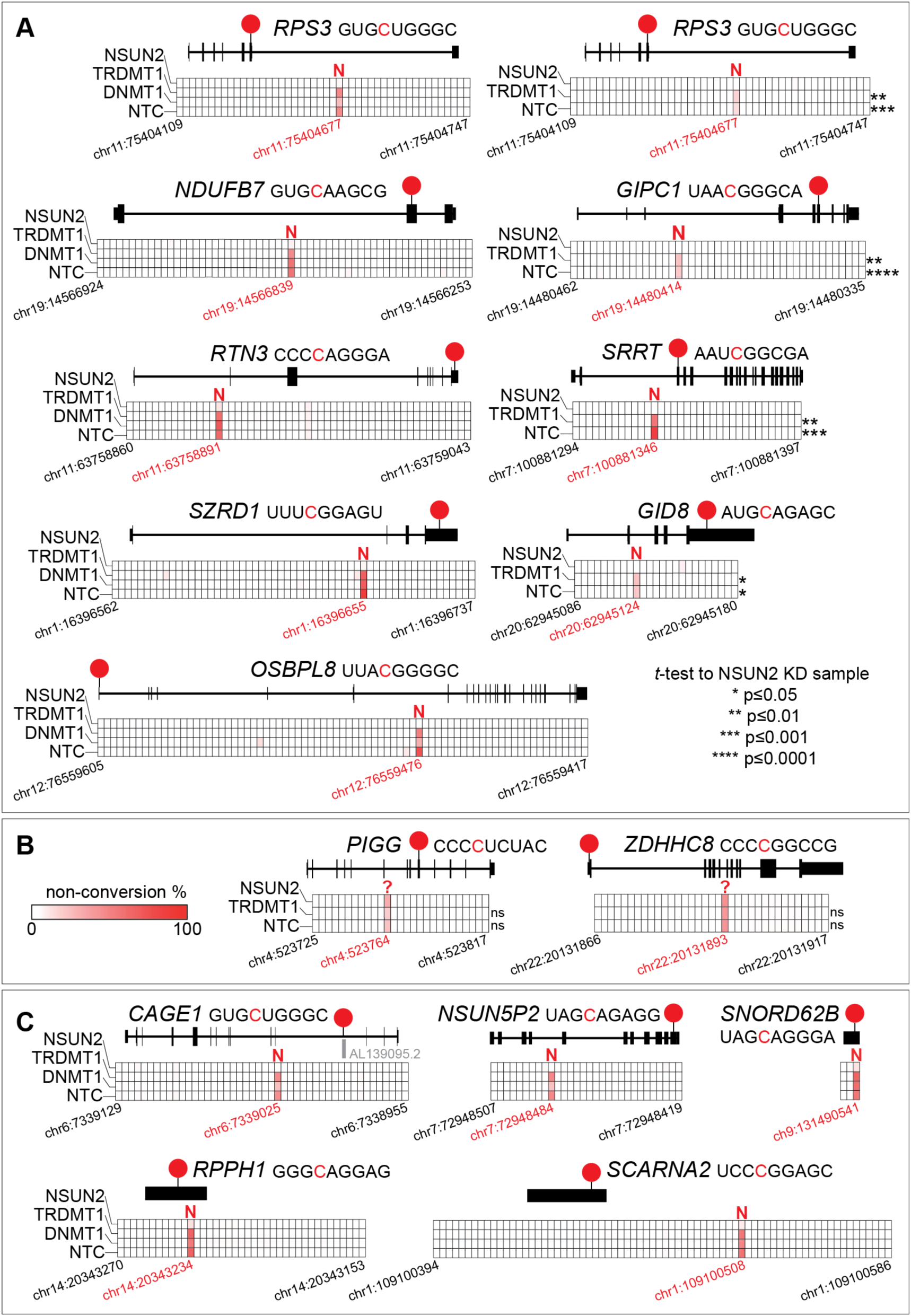
Validation and NSUN2-dependence of candidate m^5^C sites in mRNA and ncRNA. Amplicon-specific bsRNA-seq was performed with total RNA isolated from HeLa cells after siRNA-mediated m^5^C:RNA methyltransferase knockdown targeting NSUN2 or TRDMT1 along with control siRNAs (targeting m^5^C:DNA Methyltransferase 1 [DNMT1] or a non-targeting control [NTC]; see Figure S5 for knockdown efficiency controls). Read coverage achieved per amplicon was from 400 to 110,000 (see Table S6 for details). Grids in the centre of each diagram are organised by knockdown sample (in rows; siRNA target indicated on the left) and cytosine position along the analysed transcript section (in columns; genomic coordinates are given for the first and last interrogated cytosine position [in black], as well as the candidate m^5^C site [in red]). A white-to-red colour scale (shown in panel B) is used to tint each square by the degree of cytosine non-conversion observed. The longest isoform of the interrogated mRNA (based on Ensembl) is shown as a pictogram above the grid. Candidate m^5^C site position is indicated by a red circle and its sequence context given alongside the gene name (non-converted cytosine indicated in red). The enzyme identified to be responsible for cytosine methylation is indicated above the candidate sites: N – NSUN2;– unknown. ‘Confirmatory’ data (N=1) using NSUN2, TRDMT1, DNMT1 and NTC samples was generated primarily for sites with high coverage in the polysome bsRNA-seq experiment. ‘In-depth’ data (N=2 to 3) using NSUN2, TRDMT1 and NTC samples was obtained for lowly covered sites. For ‘in depth’ data, asterisks indicate p-value of paired, two-tailed Student’s *t*-test on non-conversion means between a given sample and the NSUN2 knockdown as reference (key in panel A). See Figure S6 for additional amplicon-specific bsRNA-seq analyses. A: ‘Confirmatory’ data (left panel) is shown for mRNA sites in *RPS3* (ribosomal protein S3), *NDUFB7* (NADH:ubiquinone oxidoreductase subunit B7), *RTN3* (reticulon 3), *SZRD1* (SUZ RNA binding domain containing 1) and *OSBPL8* (oxysterol binding protein-like 8). ‘In depth’ data (right panel) is shown as the average of biological replicates for mRNA sites in *RPS3*, *GIPC1* (GIPC PDZ domain family member 1), *SRRT* (serrate) and *GID8* (GID complex subunit 8 homolog). B: ‘In depth’ data for two mRNA candidate sites (*PIGG* [phosphatidylinositol glycan anchor biosynthesis class G] and *ZDHHC8* [zinc finger DHHC-type containing 8]) that showed no response to either NSUN2 or TRDMT1 knockdown. Note, the candidate site in *ZDHHC8* did not pass our coverage filter for polysome bsRNA-seq. C: ‘Confirmatory’ data for ncRNA sites in *CAGE1* (cancer Antigen 1), *NSUN5P2* (NSUN5 pseudogene 2), *SNORD62B* (small nucleolar RNA, C/D box 62B), *RPPH1* (ribonuclease P RNA component H1) and *SCARNA2* (small Cajal body-specific RNA 2). Note, the *SNORD62B* site did not have coverage in polysome bsRNA-seq.

New total RNA samples from HeLa cells as well as from two prostate cell lines (epithelial PrEC and cancerous LNCaP) were prepared to test a potential dependence of site presence on the tissue/source material (Amort et al. 2017; Yang et al. 2017). To test the MTase-dependence of sites, we used siRNA-mediated knockdown (KD) targeting NSUN2 or TRDMT1, alongside controls (siRNA targeting m^5^C:DNA Methyltransferase 1 [DNMT1] or a non-targeting control [NTC] siRNA). Knockdown efficiency was monitored for each sample by Western blotting and RT-qPCR (Figure S5A,B).

Two independent sample sets for amplicon-bsRNA-seq experiments were generated. The ‘**confirmatory**’ set (N=1) targeted sites with high cytosine non-conversion level in all three cell lines, combining HeLa cells with the full panel of siRNAs, LNCaP cells with siRNAs against NSUN2 and NTC, while PrEC cells were used in non-transfected form only. The ‘**in-depth**’ set was performed in biological triplicates (N=3) and targeted sites with lower non-conversion in HeLa cells only, combined with knockdown of NSUN2, TRDMT1 and NTC control. *In vitro* transcribed *R-Luc* spike-in controls were added prior to RNA bisulfite treatment and efficient conversion was observed for all samples (Figure S5C). Assessment of tRNA^Gly^(GCC) and tRNA^Thr^(UGU) sequences generally showed high non-conversion at the known m^5^C positions (TRDMT1 targets C38, NSUN2 targets C48-50) with complete conversion at all other assessed cytosines. Importantly, non-conversion was selectively reduced at C38 in tRNA^Gly^(GCC) in all TRDMT1 KD conditions, while the variable loop positions C48-50 strongly reacted to NSUN2 KD (Figure S5D). NTC transfection and DNMT1 KD had no such effects on tRNA non-conversion levels. Using the ‘confirmatory’ HeLa cell sample set we then confirmed three known NSUN2-dependent sites (Squires et al. 2012), in the *NAPRT* and *CINP* mRNAs (Figure S6A, top row) as well as in the ncRNA *RPPH1* (Figure 3C, bottom left; see figure legends for full gene names). Altogether this established selective MTase KD and efficient bisulfite conversion.

Next, we tested fourteen mRNA candidate m^5^C sites in HeLa cells, representing a range of coverage and non-conversion levels as well as various mRNA regions (two sites in the 5’UTR, eight in the CDS and four in the 3’UTR, respectively). Seven sites were validated in the ‘confirmatory’ samples and each showed clear reduction with NSUN2 KD (*RPS3*, *NDUFB7*, *RTN3*, *SZRD1*, *OSBPL8* in Figure 3A, left column; *SCO1*, *MCFD2* in Figure S6A, middle row). Eight sites were reproducibly detected in the in-depth samples; four showed clear and statistically significant reduction with NSUN2 KD (*RPS3*, *GIPC1*, *SRRT*, *GID8*; Figure 3A, right column). Note, that the *RPS3* site was validated with both sample sets, indicating that the ‘confirmatory’ samples are still suitable for candidate evaluation. Two sites still appeared to respond to NSUN2 KD albeit without reaching significance, likely because their low non-conversion level would require more replication (*CCT5*, *NSUN2*; Figure S6A, bottom row). Interestingly, two sites convincingly lacked responses to either MTase KD (*PIGG*, *ZDHHC8*; Figure 3B). NSUN2-independence for the *PIGG* site has been noted previously (Huang et al. 2019), we additionally show its TRDMT1-independence here. Using the ‘confirmatory’ samples, we further confirmed presence and selective sensitivity to NSUN2 KD for four sites in ncRNAs (the *RPS3 pseudogene* (AL139095.2) encoded in the *CAGE1* intron, *NSUN5P2, SNORD62B*, *SCARNA2*; Figure 3C). Finally, six candidate m^5^C sites were also explored using the two prostate cell lines (four in mRNAs: *SZRD1*, *RTN3*, *SRRT*, *PWP2*, and two in ncRNA: *SCARNA2*, *SNORD62B*; Figure S6B). All these sites were found in both PrEC and LNCaP cells and they each responded to NSUN2 KD in LNCaP cells.

In summary, the presence of all chosen sites was validated by amplicon-bsRNA-seq, even though based on polysome bsRNA-seq they varied widely in coverage (*e.g. PWP2*, *ZDHHC8*, *SNORD62B*, *NAPRT* were actually below our 30RC cut-off) and in cytosine non-conversion level. Regarding the latter, there was a reasonably good concordance between non-conversion level by transcriptome-wide and amplicon-specific measurements (e.g. *CCT5* 13% vs 5%, *RSP3* 25% vs 13% *SRRT* 63% vs 77%; averages from polysome bsRNA-seq *versus* NTC in-depth sample, respectively; see Table S6). This highlights the reliability of both, transcriptome-wide and amplicon-bsRNA-seq data. Although only based on a limited comparison, sites in both mRNA and ncRNA could be found in all three cell lines, suggesting at least some overlap in m^5^C profile between different cell types. Importantly, of the seventeen mRNA and five ncRNA sites examined by amplicon-bsRNA-seq here, all ncRNA sites and fifteen in mRNA were found to be targeted by NSUN2 and none by TRDMT1. Two mRNA sites did not respond to either knockdown and thus might be targeted by other MTases. Altogether, these data further substantiate the notion that NSUN2 has a broad but not exclusive role in modifying cellular transcriptomes (Blanco et al. 2016; Legrand et al. 2017; Yang et al. 2017; Chen et al. 2019; Huang et al. 2019; Sun et al. 2019).

### Candidate m^5^C sites display enrichment in multiple mRNA regions

∼82% of sites we identified were present in transcripts of protein-coding genes (Figure 2C). In support of specific roles, these sites are enriched for several Gene Ontology (GO) pathway terms, particularly those related to cell adhesion, translation and RNA processing/turnover (Figure S7A, Table S7). These results broadly match findings in similar cell contexts (Yang et al. 2017; Sun et al. 2019).

Given the broad role of NSUN2 in mRNA cytosine methylation, it can be expected that sites share features of the canonical tRNA substrates of the enzyme. Thus, we predicted RNA secondary structure around sites using the RNAfold tool in the ViennaRNA Package 2.0 (Lorenz et al. 2011). Compared to randomised sequences, this shows a relatively lower base-pairing tendency for the region immediately upstream of sites (position −4 to −1). Either side of this region there are patterns of alternating short segments with increased or decreased propensity for base-pairing (Figure 4A, top panel), neatly resembling the context of the major NSUN2-dependent sites in tRNA structural positions C48-50 (Figure 4A, bottom panel). We also investigated the sequence context around the modified cytosine using ggseqlogo (Wagih 2017). We noted a moderate bias for C or G in the two upstream positions and a moderate-to-strong G-bias in downstream positions 1-5, yielding a consensus of C/G-C/G-**m^5^C**-G/A-G-G-G-G (Figure 4B). Again, this consensus is similar to the C48-50 position in tRNA and its immediate 3’ sequence context. These findings elaborate on earlier reports that ‘non-tRNA’ m^5^C sites reside within CG-rich regions (Squires et al. 2012; Hussain et al. 2013a; Yang et al. 2017). The similarity to tRNA structure was further noted for sites in vtRNAs (Hussain et al. 2013a). They closely match recent findings that NSUN2-dependent sites in mRNAs reside in a sequence and structural context resembling tRNAs (Huang et al. 2019).

**Figure 4:**
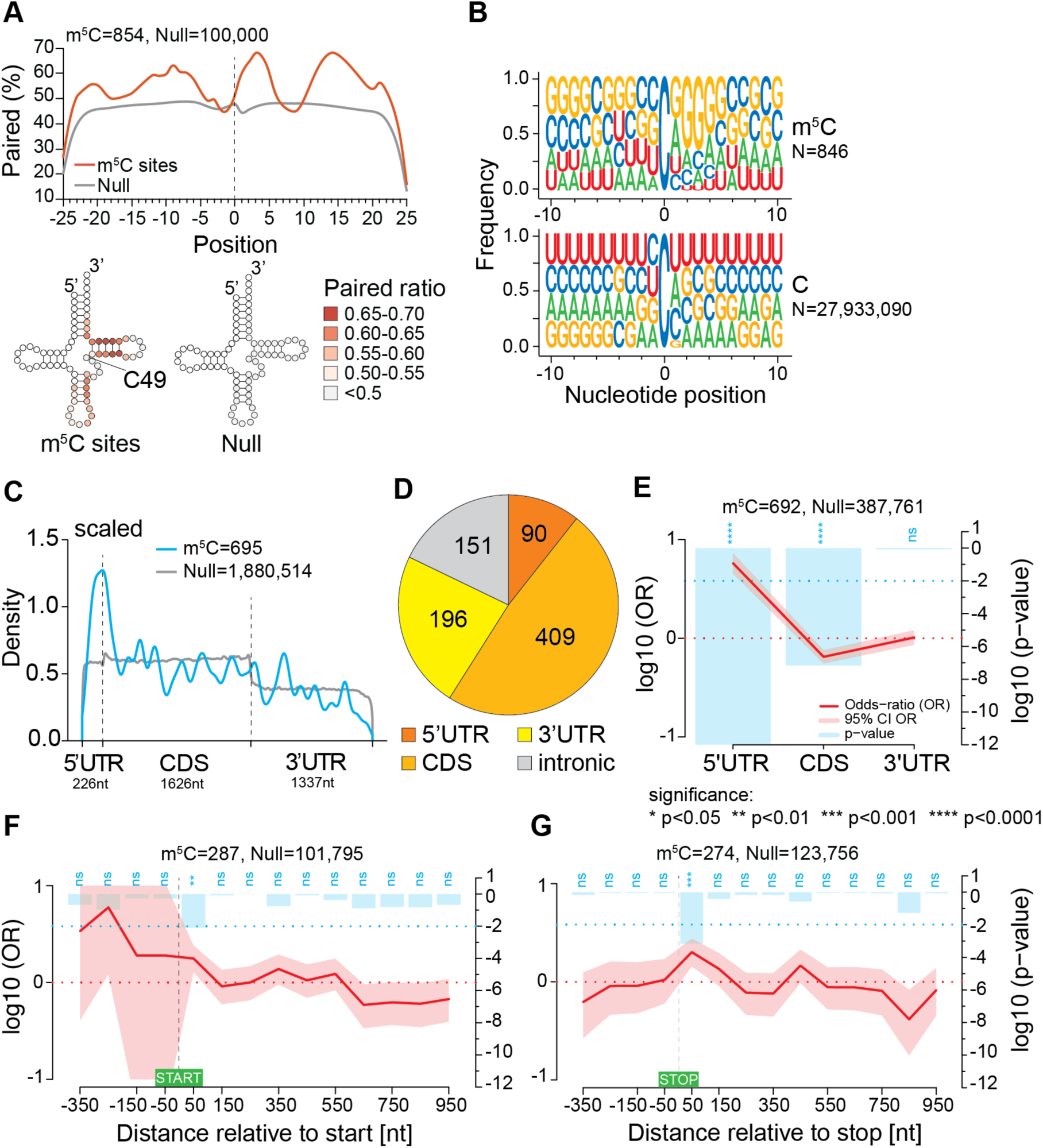
Sequence context and structural characteristics of candidate m^5^C sites. A: Base-pairing propensity meta-profile of regions surrounding candidate sites within transcripts derived from protein-coding genes (N=854). Regions around 100,000 randomly selected Cs from transcribed genome regions were used as control (top). The base pairing percentage (on a white-to-red colour scale) of regions ±20nt around candidate sites was further displayed within a cloverleaf structure, aligning candidate sites with the C49 structural position of tRNA (bottom). B: Sequence context of candidate sites in transcripts derived from protein-coding genes (N=846) (top) in comparison to all cytosines in the same transcripts (N=27,933,090) (bottom). Separate sequence logos for sites in different transcript regions (5’ untranslated region (UTR), coding sequence (CDS), 3’UTR, intronic) are shown in Figure S7C. C: Metagene density plot showing distribution of candidate sites within mature mRNAs (N=695, Null=1,880,514). Each mRNA region was scaled to its median length, indicated underneath. Candidate site distribution is shown in blue, background cytosine distribution is shown in grey. D: Pie chart showing distribution of candidate sites across 5’UTR, CDS, 3’UTR and introns of transcripts from protein-coding genes (N=846). E-G: Spatial enrichment analyses of candidate sites within mRNAs. Site are placed into chosen bins as indicated on the x-axis. Site distribution across bins is compared to matching randomised cytosine sampling (Null) and the log10 Odds-ratio (OR) is plotted as a red line with the 95% confidence interval (CI) shaded. Significance of enrichment is plotted as the log10 p-value by blue bars: ns - not significant, * <0.05, ** <0.01, *** <0.001. E: Distribution of candidate sites across the 5’UTR, CDS and 3’UTR of mRNA (m^5^C, N=692; Null=387,761). F-G: Distribution of candidate sites across the start (F) and stop (G) codon regions (from - 400nt to +1000nt relative to 1^st^ position of respective codon) using a bin width of 100nt. Site distribution across bins is compared to matching randomised cytosine sampling (Null) and the log10 Odds-ratio (OR) is plotted as a red line with the 95% confidence interval (CI) shaded. Significance of enrichment is plotted as the log10 p-value by blue bars: ns - not Note that small discrepancies in expected site numbers between panels are due to inclusion of eight sites from the ‘NMD’ RNA biotype in (A), and use of different GENCODE annotation versions, *i.e.* v28 in (A-D) and v20 in (E-G).

Next, we analysed candidate m^5^C site distribution along mRNA regions. A scaled metagene analysis showed a marked increase in sites around start codons (Figure 4C), confirming prior reports in human and mouse (Blanco et al. 2016; Amort et al. 2017; Yang et al. 2017). While sites were found in both UTRs and also in intronic regions, just under half of them were located within the CDS (Figure 4D). CDS sites showed some codon bias, being enriched in eight codons specifying 5 amino acids, primarily in the first and second codon positions (Figure S7B). Despite the numerical predominance of CDS sites, spatial enrichment analysis of sites in mRNA using RNAModR (Evers et al. 2016) revealed significant overrepresentation of sites in the 5’UTR, with a minor but significant underrepresentation in the CDS (Figure 4E). Of note, adherence to the site sequence context established above (Figure 4B) was strongest in the 5’UTR, with some divergence in the 3’ UTR (consensus: G/C-U/G-**m^5^C**-A/G-G-G-G-G (Figure S7C). To further inspect site prevalence near start codons, we divided the surrounding region (−400 to +1000nt) into 100nt bins and directly tested for enrichment (Figure 4F). The broad window and relatively coarse bin size were necessary to retain statistical power, given the relatively low site numbers in mRNA. This showed a gradient of decreasing site prevalence in 5’ to 3’ direction, confirming the concentration of sites along 5’UTRs. Within 5’UTRs, the −201 to −300 interval showed the highest odds ratio, albeit without reaching significance. As the median length of 5’UTRs represented in our data is 226nt, this could suggest some concentration of candidate m^5^C sites near the mRNA 5’ end, however, site numbers are too low to ascertain this (Figure S7D, left panel). The 100nt region immediately downstream of the start codon showed significant site enrichment, while bins within the body of the CDS (> 600nt downstream of start codons) showed a continued decrease of site density (Figure 4F). We extended these analyses to several other mRNA features, which mostly remained inconclusive due to diminishing site numbers in any given region. There was, however, a significant site enrichment within the 100nt interval immediately downstream of stop codons (Figure 4G) and in the interval 101-200nt upstream of mRNA 3’ ends as well (Figure S7D, right panel). Altogether, the diversity of candidate m^5^C site distribution patterns observed here hint at distinct, context-dependent functional roles for m^5^C.

### Transcriptome-wide anti-correlation between cytosine modification level and mRNA translation efficiency

Links to mRNA translation are suggested by several of the site enrichment patterns described above. This was also emphasised in a recent report, showing a significant negative correlation between candidate m^5^C site-content in the CDS and translation efficiency in the HeLa cell transcriptome (Huang et al. 2019). To independently verify this, we obtained HeLa cell ribosome profiling data from several studies (Wang et al. 2014; Park et al. 2016; Arango et al. 2018). Cumulative distribution of translation efficiency values allowed us to compare mRNAs found by us to contain candidate m^5^C sites with the remaining mRNAs. Irrespective of underlying ribosome profiling data, we then saw a clear and significant tendency for site-containing mRNAs to be less well translated (Figure S8A). In a similar vein, we also assessed any transcriptome-wide relationship with mRNA stability (Wang et al. 2014; Arango et al. 2018), and found site-containing mRNAs to display a significant trend towards longer half-life (Figure S8B). Interestingly, this latter observation matches findings reported recently for m^5^C-modified mRNAs in mammals and zebrafish (Chen et al. 2019; Yang et al. 2019b).

A unique advantage of our sampling approach is that it allows us to profile the level of cytosine non-conversion at each site across polysome gradient fractions, with bsRNA-seq data for each fraction available in biological triplicates (see Figure S2, Table S8A). We first assessed the overall non-conversion range of the 846 sites in transcripts from protein-coding genes in each fraction. This showed declining non-conversion levels with increasing ribosome association, with comparisons of fraction 1-to-2 and 2-to-3 reaching statistical significance (Figure 5A, left panel). This trend was most pronounced with sites in the CDS and still discernible with 5’UTR sites, whereas the 3’UTR and intronic sites did not show a clear trend (Figure 5A, remaining panels). This demonstrates a negative correlation, on the bulk level, between the extent of cytosine non-conversion and mRNA translation state, primarily driven by observations with CDS sites.

**Figure 5:**
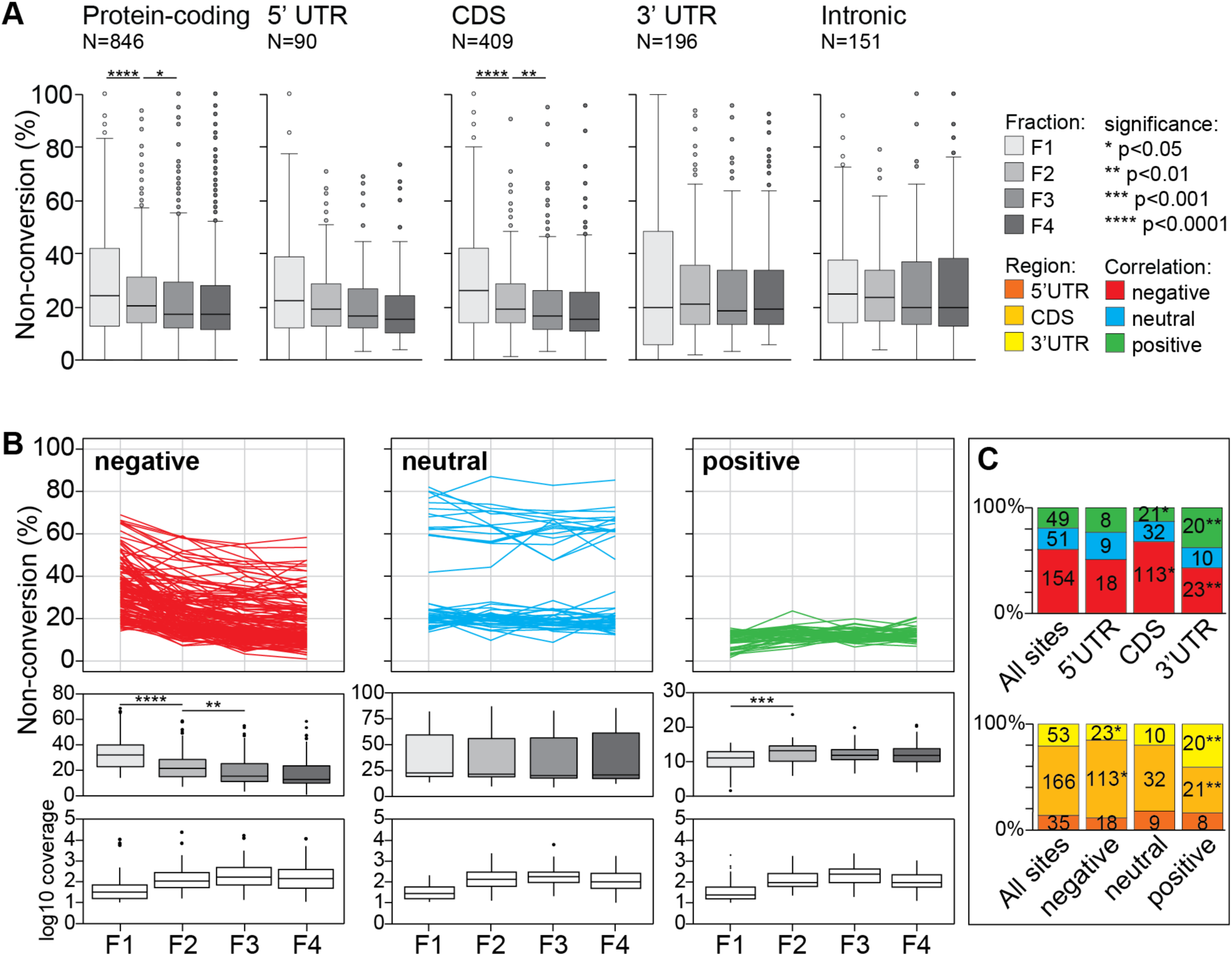
Relationship between non-conversion level at candidate m^5^C sites and mRNA translation state. bsRNA-seq libraries were grouped as biological triplicates per fraction (*e.g.* LibB1, LibC1 and LibE1 each report on sites detected in bsRNA-seq fraction 1), allowing the calculation of average non-conversion levels per individual site and per fraction. A: Boxplots showing distribution of candidate site non-conversion levels across the polysome profile. Shown from left to right are: all sites in protein-coding RNA (N=846, *c.f.* Figures 2C,4D), as well as subsets of these sites in the 5’ untranslated region (UTR, N=90), coding sequence (CDS, N=409), 3’UTR (N=196) and intronic (N=151). Asterisks indicate significance p-value from unpaired, two-tailed Student’s *t*-test comparing the means of adjacent fractions. B: Non-conversion levels per individual site across the polysome profile were partitioned into nine soft clusters using Mfuzz (see Figure S9). 254 candidate m^5^C sites in exonic mRNA regions (5’UTR, CDS, 3’UTR designation in panel A) were included based on having coverage in at least 9 out of 12 bsRNA-seq fraction samples and ≥10 average read coverage in each of the four bsRNA-seq fractions. Mfuzz clusters were grouped into three translation state trend categories by visual inspection, showing a negative (N=154), neutral (N=51) or positive trend (N=49) with polysome association. Top panels: line graphs displaying individual site average non-conversion levels across fractions. Middle panels: boxplots showing distribution of site non-conversion levels in each fraction. Asterisks indicate significance p-value from unpaired, two-tailed Student’s *t*-test comparing the means of adjacent fractions. Bottom panels: boxplots showing distribution of site read coverage in each fraction. C: Stacked bar charts showing distribution of sites in the different translation state trend categories across mRNA regions (top) and distribution of sites in different mRNA regions across translation state trend categories (bottom). Asterisks indicate significance p-values following binomial test against the distribution of all sites. The legend to the right of panel A gives a colour/significance key applicable to all panels. See Figure S10 for cluster analysis with additional sites with sufficient coverage only in bsRNA-seq fractions 2-4.

Next, we considered non-conversion levels of sites individually and performed Mfuzz clustering (Kumar and M 2007). We selected a set of sites in mature mRNA that had sufficient (≥10 reads average) coverage in all 4 fractions (F1234; 254 sites), as well as a second set that had sufficient coverage in fractions 2-4 but not in fraction 1 (F234; 315 sites). Mfuzz was run requiring 9 clusters (Figures S9,S10A, Table S8B,C) before re-grouping clusters based on overall non-conversion trends. This generated three major profile patterns for each set, representing positive, neutral and negative correlation with translation state, respectively. Positive and negative pattern sets each showed significant non-conversion level change between fractions as expected (F1234 shown in Figure 5B; F234 shown in Figure S10B). Focusing on the set with stronger discriminative potential, F1234, we saw that non-conversion levels were not distributed across patterns equally; most notably, sites with positive patterns typically had low non-conversion levels through the fractions, whereas sites with a neutral pattern were, for unknown reasons, split into two groups, one with ∼20% and a smaller group with ∼60% non-conversion (Figure 5B, top panels).

Notably, site profiles indicating negative correlation were the most common (∼61%), with positive profiles (∼19%) being the least frequent (Figure 5C, top panel). This bias towards negative profiles was moderately but significantly enhanced with CDS sites (∼68%), whereas 3’UTR sites were significantly underrepresented (∼43%). Among profiles showing positive correlation with translation, CDS sites were significantly depleted (∼13%), while 3’UTR sites (∼38%) were significantly enriched. Conversely, compared to all sites, those in the negative pattern set were moderately but significantly enriched for CDS location and depleted for 3’UTR location. Sites with a positive pattern were depleted for CDS location but enriched for 3’UTR location (Figure 5C, bottom panel). Many, but not all, of these observations were also made in the less discriminative F234 set (Figure S10). With the caveat that a proportion of individual site profiles are based on imprecise measurements (see below), the key discernible features from the clustering approach were a) a preponderance of sites showing negative correlation with translation state, and b) while 5’UTR sites were relatively unremarkable, there was a tendency for CDS and 3’UTR site to segregate into negative and positive pattern sets, respectively.

### Individual mRNA sites show robust anti-correlation with translation state

To identify individual sites displaying significant non-conversion change across the polysome profile, we performed pairwise logistic regression analysis (Table S8D). In each pairwise comparison, the majority of sites that reached significance showed a negative correlation with translation state; the F1-F2 comparison yielded the largest number of significant sites with the strongest bias towards the negative trend (Figure S11A). 43% of the sites in the F1234 set but only ∼7% in the F234 set, had significant differences in at least one pairwise comparison (Table S6). Focusing on the F1234 set, 108 sites reached significance comprising a total of 149 significant pair-wise comparisons. Of note, the large majority of these sites represented a negative trend with translation state, and most assignments were based on the F1-F2 and F2-F3 comparisons (Figure S11B). Sixteen of these sites were in the 5’UTR, 81 in the CDS, and 11 in the 3’UTR, which represents significant enrichment of CDS sites and depletion of 3’UTR sites.

Sites with significant change in cytosine non-conversion level across several fraction steps, or with larger magnitude of change between steps, may represent the most compelling candidates for functional studies. Regarding the former, 12 sites reached significance in all three pairwise comparisons; 10 of these showed a negative trend (17/79 with 14/69 sites showing a negative trend based on two, or a single pairwise comparison, respectively). Regarding the latter, 59 of the 149 significant pairwise comparisons satisfied an arbitrarily imposed criterion of ≥10% relative non-conversion difference. 54 of these pairs represented a negative step. Furthermore, they were primarily based on the F1-F2 comparison (39; 15 on F2-F3 and 5 on F3-F4). In terms of sites, none of the twelve ‘triple significance’ sites satisfied this criterion at all steps, one having two steps and five having one step of the required magnitude (for the 17 ‘double significance’ sites: three for two steps, twelve for one step; 79 ‘single significance’ sites: 33 for one step). Overall, choosing sites for ‘compelling’ profiles across the polysome gradient primarily, but not exclusively, selects for those showing a negative trend with translation state.

The profiles of sites selected for significant (for two or more steps) and/or strong change (≥10% relative), as well as different level of cytosine non-conversion overall, are shown in Figure 6 and Figure S11C,D. The selection further contains several sites that were validated by amplicon-bsRNA-seq. Inspecting the few ‘promising’ sites with positive change showed that several of them actually displayed a complex profile pattern, consistent with their varied membership to the trend patterns described above (Figure 5B) and leaving even fewer with a clear, monotonously positive association with translation state (Figure S11D). By contrast, all ‘high quality’ sites with negative change came from the negative trend pattern (Figure 5B, left panel) and nearly all displayed a continuous decline in cytosine non-conversion from fraction 1 through to 4 (Figures 6,S11C). While a few of these latter sites were situated in the 5’ or 3’UTR, most of them were located in the CDS. Thus, bsRNA-seq has identified candidate m^5^C site in multiple individual mRNAs that suggest an interdependence with translation. These sites/mRNAs are now accessible to functional follow-up studies.

**Figure 6:**
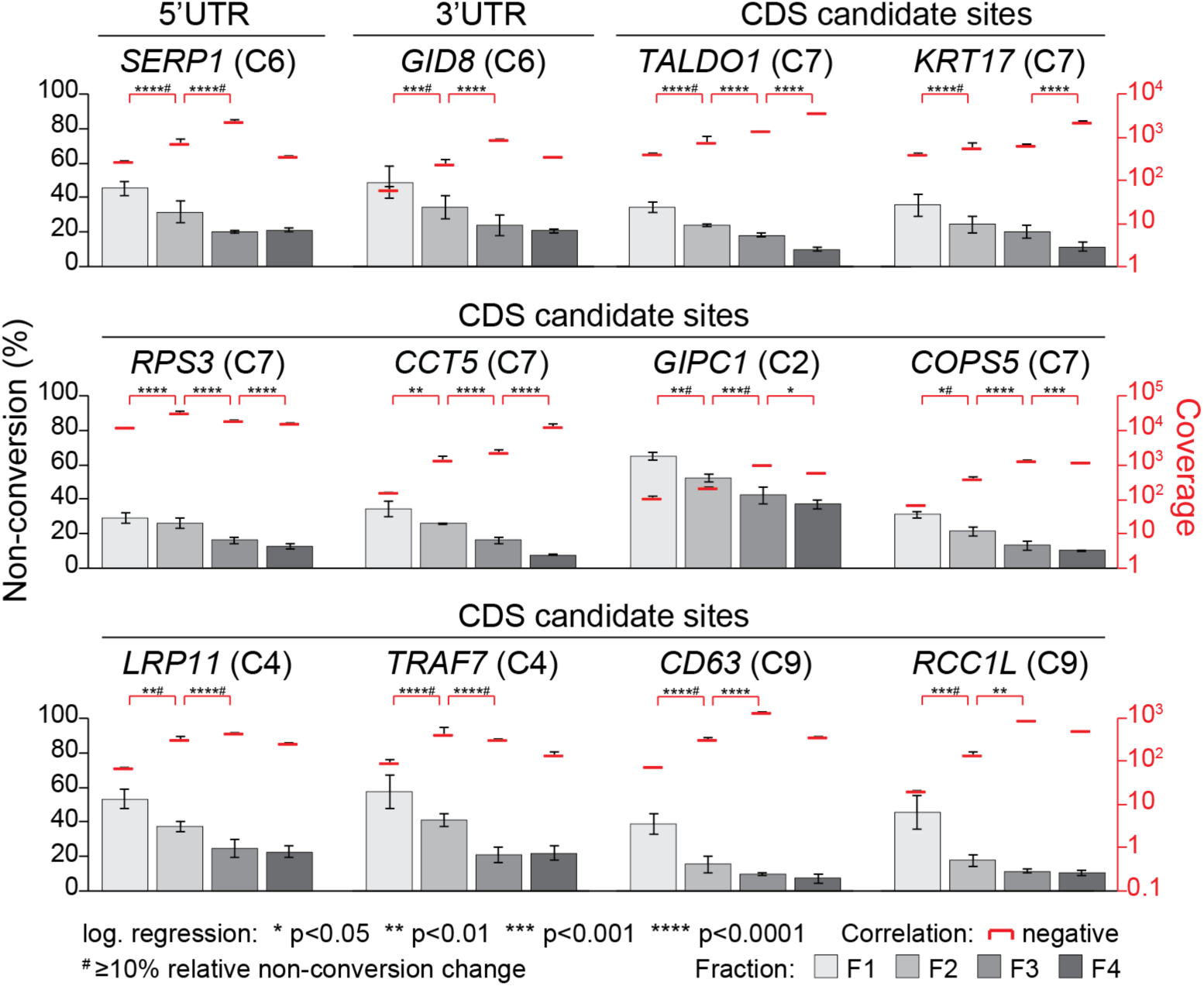
Examples of individual candidate m^5^C sites in mRNA showing significant correlation of cytosine non-conversion with translation state. Dual axis charts show cytosine non-conversion (bars) and coverage (red lines) for a given site across the polysome gradient. Data is shown as means across biological triplicates, with error bars indicating standard deviation. Asterisks indicate significance p-values after logistic regression testing (see key below the charts). The gene name for each candidate site and its position within the mRNA is given. All sites shown here show significant negative non-conversion change in at least two fraction steps and are represented in the F1234 clustering with the cluster number indicated in brackets. mRNAs shown are: *SERP1* (stress associated endoplasmic reticulum protein 1); *GID8* (GID complex subunit 8 homolog); *TALDO1* (transaldolase 1); *KRT17* (keratin 17); *RPS3* (ribosomal protein S3); *CCT5* (chaperonin containing TCP1 subunit 5); *GIPC1* (GIPC PDZ domain containing family member 1); *COPS5* (COP9 signalosome subunit 1); *LRP11* (LDL receptor related protein 11); *TRAF7* (TNF receptor associated factor 7); *CD63* (CD63 molecule); *RCC1L* (RCC1-like)

## Discussion

We present here a set of ∼1,000 high-confidence candidate m^5^C sites in the human HeLa cell transcriptome. The great majority of sites were found in mRNAs, and their sequence and structure contexts strongly resembled that of the canonical NSUN2 target region in the variable loop of tRNAs. This matches our extended validations by amplicon-bsRNA-seq, which attributed 21 of 23 confirmed sites to NSUN2. Several findings point towards functional links particularly to translation, including site enrichment in the mRNA 5’ region and near start codons. m^5^C-containing mRNA species further display relatively lower translation efficiency as measured by ribosome profiling. Uniquely, we exploited the generally sub-stochiometric modification level of mRNAs to directly show a prevailing negative correlation between modification state and recruitment into polysomes.

The merits of using bsRNA-seq to discover m^5^C sites transcriptome-wide have been controversially discussed (Legrand et al. 2017; Linder and Jaffrey 2019; Motorin and Helm 2019; Trixl and Lusser 2019). We contend that the bsRNA-seq approach as presented here is fit-for-purpose as it is based on a combination of efficient bisulfite reaction conditions, bespoke read mapping and conservative site calling from replicate data. Nevertheless, our operationally defined settings to reduce false positives also incur limitations. As illustrated for tRNAs, the ‘3C’ criterion biases against detection of more than three closely-spaced sites. The ‘S/N90’ criterion hinders detection of true sites within strong RNA secondary structure, as shown for rRNA. Still, these setting are justifiable given that our focus was on sites elsewhere in the transcriptome. Similarly, applying the ‘10MM’ criterion to remove sites with low non-conversion level seems prudent to focus on functionally important sites. However, the use of thresholded data would be problematic for comparative studies of methylation level between tissues or treatment conditions. It may also be unnecessarily stringent when using the data for other purposes, for example when characterising MTase substrate requirements. Supporting the overall validity of our approach is our high amplicon-based validation rate and the extensive site overlap with a recent study that converged on a similarly stringent approach to analyse bsRNA-seq data (Huang et al. 2019).

Short of an unexpected discovery of novel RNA:m^5^C MTases, sites in the eukaryotic transcriptome at large need to be deposited by one of the existing NSUN enzymes or TRDMT1. The well-characterised NSUN2 has long been the prime suspect, and there is indeed cumulative evidence pointing to NSUN2 modifying both, mRNA and ncRNA from earlier studies (Squires et al. 2012; Hussain et al. 2013b; Khoddami and Cairns 2013; Blanco et al. 2016; Yang et al. 2017; Huang et al. 2019; Sun et al. 2019). We examined this here by amplicon-bsRNA-seq and found 5/5 ncRNA sites and 15/17 mRNA sites tested to be targeted by NSUN2. None was targeted by TRDMT1, leaving two mRNA sites that might be targeted by other MTases. This independently confirms, and adds to, earlier targeted analyses of this kind. For example, the existence and NSUN2-dependence of sites in the ncRNAs *RPPH1*, *SCARNA2* and several vault RNAs has been repeatedly shown (Squires et al. 2012; Hussain et al. 2013b; Khoddami and Cairns 2013). Two of the fifteen mRNA sites shown to be NSUN2-dependent here, were reported by us before (Squires et al. 2012). Thirteen mouse mRNA sites discovered by bsRNA-seq were independently validated by m^5^C-RIP-seq, although their NSUN2-dependence was not assessed (Amort et al. 2017). Further, we and others (Huang et al. 2019) have found that, transcriptome-wide, m^5^C site context resembles that of the major NSUN2-dependent sites in tRNA structural positions C48-50. Thus, NSUN2 appears to identify all of its targets by recognising tRNA-like features. Curiously, while in tRNA two or three adjacent cytosine are often modified by NSUN2, this seems to be rare in its non-canonical targets. It was also found that most, but not all, transcriptome-wide sites lacked methylation in NSUN2 knock-out cells; those that were NSUN2-independent were situated in a distinct sequence/structural context (Huang et al. 2019). Related to that, it was very recently reported that a subset of human mRNA m^5^C sites showed differential cytosine modification levels in TRDMT1 knockdown cells; one site was independently validated (Xue et al. 2019). Altogether, multiple lines of evidence implicate NSUN2 as a major, but likely not the only, MTase targeting mRNAs.

Available information on epitranscriptomic marks suggests a range of context-dependent functions and m^5^C is likely no exception. Several observations in this study individually, but not necessarily coherently, suggest functional links to mRNA translation but also stability, broadly concurring with prior evidence. Drawing on published mRNA half-life and ribosome profiling data we saw that, compared to the rest of the transcriptome, m^5^C-containing mRNAs were more stable but less-well translated. A negative transcriptome-wide correlation between m^5^C site presence in human mRNAs and mRNA translation efficiency, as measured by ribosome profiling, was shown before (Huang et al. 2019) as was a positive transcriptome-wide association with mammalian and zebrafish mRNA stability (Chen et al. 2019; Yang et al. 2019b). Further, effects of m^5^C site on stability and translation of several individual mRNAs in the context of cell senescence have been reported (Wang 2016; Casella et al. 2019). Given that these are opposing trends in terms of gene expression output, a reasonable guess would be that different subsets of sites drive each association. Uniquely, by comparing polysome-association of modified with unmodified mRNAs of the same type, we directly showed anti-correlation between cytosine modification level and translation efficiency. This was true for most, but not all mRNAs available for this analysis, again suggesting diversity of site functionality. Distribution of candidate m^5^C sites was also uneven in mRNAs. We saw a gradient of increasing site prevalence towards the mRNA 5’ end, resulting in an enrichment in the 5’UTR. Overall, CDS sites showed enrichment in the first and second positions of eight codons. We could also discern site enrichment just downstream of start and stop codons as well as upstream of the mRNA 3’ end. Enrichment of m^5^C sites around the start codon region has been noted repeatedly (Blanco et al. 2016; Amort et al. 2017; Yang et al. 2017; Chen et al. 2019). Site positioning could be a key determinant of downstream function, and thus might co-segregate with other patterns suggesting distinct functionality. Unfortunately, for the most part we lacked sufficient site numbers to conduct meaningful statistical analyses of this kind. Nevertheless, we did see an enrichment of CDS sites and a depletion of 3’UTR sites among mRNAs showing a negative correlation with translation.

Demonstrating and mechanistically characterising specific causal links between m^5^C and mRNA translation will require in-depth study of prototypical examples; another contribution of this study is that we provide multiple such mRNAs that can be analysed in future work.

## Methods

### Cell growth and maintenance

HeLa (Human Cervical Cancer) cells were obtained from ATCC and grown in DMEM medium (Life Technologies) supplemented with 5% FBS and 2mM L-glutamine and passaged when 70-100% confluent. Prostate cell lines were obtained from ATCC. PrEC cells (Prostate Epithelial Cells) were cultured in PrEBM basal medium supplemented with PrEGM SingleQuots™ Supplements and Growth Factors (Lonza) and passaged when 80-90% confluent. LNCaP cells (Prostate Cancer Cell) were cultured in T-Medium (Life Technologies) supplemented with 10% FBS and 2mM L-glutamine and passaged when 90-100% confluent. All cells were incubated at 37°C with 5% CO_2_.

### Generation of the R-Luc spike-in control RNA sequences

Humanised Renilla Luciferase (*R-Luc*) RNA, transcribed *in vitro* from the *R-Luc* insert located in pCl-Neo (Bhattacharyya et al. 2006), was used as a bsRNA-seq spike-in negative control to assess conversion efficiency. Either this RNA or a second non-humanised *R-Luc* RNA, transcribed *in vitro* from the *R-Luc* insert in the pRL-TK vector, was used as negative control in the amplicon-bsRNA-seq experiments. RNAs were transcribed using the MEGAscript® T7 Kit (Life Technologies) according to the manufacturer’s protocol. Template DNA was removed using TURBO™ DNase (Ambion) and the RNA cleaned by phenol/chloroform extraction and precipitated. A second DNase treatment step was performed to remove any residual template DNA. The size and integrity of each *in vitro* transcript was assessed using the RNA 6000 Nano Kit on the 2100 Bioanalyzer (Agilent Technologies).

### Western blotting

For Western blotting of sucrose gradient fractions, the protein fraction samples were used directly for gel electrophoresis without any further protein purification. For Western blotting of methyltransferase knockdown samples, protein was isolated from cells using 300μl CytoBuster™ Protein Extraction Reagent (Novagen) per well according to manufacturer’s instructions. Western blotting was performed as standard, separating proteins on NuPage™ 4-12% Bis-Tris Protein Gels (Invitrogen) followed by transfer onto nitrocellulose or PVDF (for IR-Dye detection) membrane. The membrane was blocked in 5% milk in PBS-T (0.05% Tween-20) or Odyssey Blocking Buffer (for IR-Dye detection; LI-COR 927-40000) and probed with primary antibodies: anti-RPL26 antibody (1:1,000; Sigma R0780), anti-NSUN2 (1:1,000 – 1:5,000; Proteintech 20854-1-AP), anti-TRDMT1 (1:1,000; Proteintech 19221-1-AP), anti-tubulin (1:1,000; Sigma T6199) or anti-GAPDH (1:1,000; Abcam ab9484) at room temperature. The membranes were probed with a secondary antibody: either anti-rabbit-HRP (1:5,000; Merck Millipore AP132P), anti-mouse-HRP (1:5,000; Agilent Dako P0260), anti-mouse-IR-Dye800 (1:10,000; LI-COR 926-32210) or anti-rabbit-IR-Dye680 (1:10,000; LI-COR 925-68071). For HRP detection, membranes were incubated with substrate (Pierce) and visualised. For IR-Dye detection, the membranes were scanned using the Odyssey® CLx Imaging System (LI-COR).

### Amplicon bisulfite RNA sequencing data generation and analysis

#### siRNA-mediated knockdown of methyltransferase genes

Gene knockdown was performed by Lipofectamine® RNAiMax (Invitrogen) transfection with SMARTpool siGENOME siRNAs (Dharmacon) targeting *NSUN2*, *TRDMT1*, *DNMT1* and a non-targeting control (NTC) as described previously (Squires et al. 2012). Briefly, 1.5 x 10^5^ cells were transfected with 60pmol siRNAs in a six-well plate format, passaged after 3 days, transfected again and harvested 6 days post initial transfection. To assess knockdown efficiency, protein and RNA were isolated for Western blot (as described above) and reverse transcription followed by quantitative PCR (RT-qPCR) analyses, respectively. Of note: NSUN2 and NTC siRNA transfection of prostate cells was only performed in LNCaP cells as PrEC cells showed strongly decreased viability in response to transfection.

#### RNA extraction and reverse transcription-quantitative PCR (RT-qPCR)

RNA was extracted with TRIzol (Invitrogen) or TRI Reagent® (Bian et al.) according to manufacturer’s instructions. Briefly, cells were suspended in 1ml TRI Reagent per well, mixed with 200μl chloroform and incubated for 3min. Phases were separated by centrifugation at 12,000rpm for 15min at 4°C. RNA was precipitated from the aqueous phase for 10min with 500μl isopropanol in the presence of 5μl glycogen (10mg/ml). RNA was collected by centrifugation at 12,000rpm for 10min at 4°C. The pellet was washed with 75% ethanol, suspended in nuclease-free water and ethanol precipitated overnight at −20°C. Precipitates were collected by centrifugation at 12,000rpm for 15min at 4°C, the pellets washed with 75% ethanol and resuspended in nuclease-free water. RNA samples were analysed for quality using a Thermo Scientific™ NanoDrop™ One spectrophotometer (Thermo Fisher) and for integrity using the RNA 6000 Nano Kit on the 2100 Bioanalyzer (Agilent Technologies). RNA was treated using the TURBO DNA-*free*™ Kit (Invitrogen) according to manufacturer’s instructions, purified by phenol/chloroform extraction and precipitated. An RNA subsample was reverse-transcribed using SuperScript™ III Reverse Transcriptase (Invitrogen) with oligo-dT primers and qPCR performed using Fast SYBR^®^ Green Master Mix (Applied Biosystems). Primer sequences are listed in Table S2. Amplifications were performed in technical triplicates in a 384-well format using the QuantStudio™ 12K Flex system (Life Technologies). Housekeeping genes (e.g. *GAPDH* or *ACTB* as indicated) were used as internal standard control to normalise expression levels.

#### Bisulfite conversion and amplicon-bsRNA sequencing

The remaining RNA was spiked with 1/1000 (w/w) of *R-Luc in vitro* transcript and 1μg of RNA subjected to bisulfite conversion as described previously (Sibbritt et al. (2016). Purified RNA was reverse-transcribed using SuperScript™ III Reverse Transcriptase (Invitrogen) with random hexamers and subjected to amplicon-specific touchdown PCR. PCR conditions were optimised for each primer set and carried out in technical triplicates (primer sequences specifically target bisulfite converted templates and are shown in Table S3). The obtained products were analysed by agarose gel electrophoresis, purified using the Wizard^®^ SV Gel and PCR Clean-Up System (Promega) or MinElute^®^ PCR Purification Kit (Qiagen) and the technical replicates combined. PCR products from the same RNA sample were pooled and subjected to library preparation. ‘Confirmatory’ libraries were prepared using the TruSeq^®^ DNA LT Sample Prep Kit with minor modifications, whereas ‘in depth’ libraries were prepared using the TruSeq^®^ DNA Nano Library Kit (Illumina) according to manufacturer’s instructions. Libraries were mixed with 50% PhiX Control Library (Illumina) and sequenced on the MiSeq System (Illumina), acquiring 151bp read length. Of note: For the ‘confirmatory’ HeLa and Prostate datasets only a single biological replicate for each sample was analysed, except for PrEC and LNCaP wild-type samples, which were performed in biological duplicates. For the ‘in depth’ HeLa dataset, all analyses were performed in biological triplicates.

#### Mapping of amplicon-bsRNA-seq reads and site calling

The target mRNA regions of all amplicons were combined into a single reference and the forward strand C-to-T converted. Reads from ‘confirmatory’ libraries were first trimmed using Trimmomatic (Bolger et al. 2014) in palindromic mode with parameters (ILLUMINACLIP:illuminaClipping.fa:4:30:10:1:true LEADING:3 TRAILING:15 SLIDINGWINDOW:4:15 MINLEN:36). Sequencing reads were aligned using Bowtie2 within Bismark (Krueger and Andrews 2011). Reads from ‘in depth’ sequencing were subjected to FastQC (v0.11.5); adapter removal and low-quality base trimming was performed using cutadapt (v1.18) with options (-q 20,20 -m 50 --trim-n -a AGATCGGAAGAGCACACGTCTGAACTCCAGTCA –A AGATCGGAAGAGCGTCGTGTAGGGAAAGAGTGT). Clean reads were aligned to the reference using the MeRanT tool in meRanTK (Rieder et al. 2016) in both directions, as sequencing was not strand-specific, and the resulting bam files merged. Non-conversion level was determined for each C position within the amplicons.

### Polysome bisulfite RNA sequencing data generation and analysis

#### Sucrose density gradient centrifugation and polysome profiling

For polysome profiling, HeLa cells were grown until ∼70% confluent and then 2 technical replicate 150mm diameter plates were seeded with 6 x 10^6^ cells per biological replicate. Cells were again grown to ∼70% confluency, and lysates prepared in the presence of cycloheximide (200μl lysis buffer per plate), essentially as previously described (Clancy et al. 2007). The lysate protein concentrations were measured by *DC* protein assay (Bio-Rad) according to manufacturer’s instructions and by absorption at 260nm. Lysates were frozen and stored at −80°C until further processing. To recover the maximum amount of RNA possible, the entire lysate volume (∼500μl) was layered on top of a 10ml 17.5-50% linear sucrose density gradient tube and separated by ultracentrifugation. Then, fractions were collected from the top of the gradient using a Brandel Gradient Fractionator (flow rate of 0.75ml/min), collecting 24 fractions at 36s intervals while measuring the absorption at 254nm. 10μl subsamples were removed from each of the 24 fractions and combined by three to obtain a total of 8 protein fractions for downstream protein distribution analysis by Western blotting (as described above). All 24 collected fractions (0.5ml) were spiked-in with 1ng *in vitro* humanised *R-Luc* transcript to control for variations in RNA isolation efficacy. RNA was precipitated from each fraction with 1.5ml ethanol in the presence of 5μl glycogen (10mg/ml) at −80°C overnight before proceeding to RNA extraction. RNA samples were combined by two to obtain 12 RNA fractions and subjected to a second ethanol precipitation step to remove residual sucrose. Yield and integrity of the isolated RNA was assessed using the RNA 6000 Nano Kit on the 2100 Bioanalyzer (Agilent Technologies). Three independent biological replicates were prepared in this way, with at least one cell passage between each. Key parameters for each replicate can be found in Table S1.

#### Reverse transcription and quantitative PCR (RT-qPCR)

DNA contamination was removed from RNA using the TURBO DNA-*free*™ Kit (Invitrogen) according to manufacturer’s instructions. RNA was purified by phenol/chloroform extraction and precipitated. A subsample from each RNA fraction was reverse-transcribed using SuperScript™ III Reverse Transcriptase (Invitrogen) with oligo-dT primers and qPCR amplifications were performed in technical triplicates using Fast SYBR^®^ Green Master Mix (Applied Biosystems) in a 384-well format using the QuantStudio™ 12K Flex system (Life Technologies). Primer sequences are listed in Table S3. The *in vitro R-Luc* spike-in transcript was used as control for variation in RNA isolation efficiency between each fraction and all results are represented relative to *R-Luc*.

#### Bisulfite conversion and bsRNA-seq

The remaining RNA was combined into four final bsRNA-seq fractions (Figure 1) and 10μg RNA was spiked with ERCC Spike-in Mix 2 (Ambion) according to manufacturer’s instructions. RNA was then used for sequencing library construction using the TruSeq^®^ Stranded Total RNA Library Kit (Illumina) according to manufacturer’s instruction, with some modifications. Following rRNA depletion, RNA was suspended in nuclease-free water and subjected to bisulfite conversion as described previously (Sibbritt et al. (2016). Converted RNA was purified, and 1μg used for continued library preparation with omission of the fragmentation step as the RNA undergoes fragmentation during the bisulfite treatment. No size-selection was performed as the size of the bisulfite treated RNA fragments was between 50-200nt with a peak size of approximately 150nt (Figure S1). The libraries were mixed equally and loaded onto the HiSeq 2500 System (Illumina) using a total of three lanes for sequencing. Sequencing was performed in fast mode acquiring 101bp paired-end reads.

#### Preparation of reference sequences

The human reference genome hg38 and the GENCODE v28 annotation for each chromosome were downloaded from UCSC. The human ribosomal DNA complete repeating unit (U13369.1) was downloaded from NCBI and the human hg38 tRNA annotations were downloaded from GtRNAdb (Chan and Lowe 2016). To assemble a pre-tRNA reference, 3’ and 5’ genomic flanking regions of length 100nt with the corresponding tRNA reference were extracted from the genome with BEDTools. Intronic sequences were also included. ERCC spike-in reference sequences (SRM374) were obtained from www-s.nist.gov, the *R-Luc* sequence is listed in Table S2. The reference genome sequence as well as ERCC and *R-Luc* spike-in sequences were combined into a single reference sequence. Then converted as follows: C-to-T conversion of the forward strand and G-to-A conversion of the reverse strand followed by indexing using ‘meRanG mkbsidx’. tRNA and rRNA sequences were each treated as separate references and only C-to-T converted followed by indexing using ‘meRanT mkbsidx’ (Figure S2).

#### Initial read mapping of bsRNA-seq reads

Raw reads were subjected to FastQC (v0.11.5). Low quality bases and adaptor sequences were removed using Trimmomatic (v0.36, (Bolger et al. 2014)) with options (ILLUMINCLIP:Adapter.fa:2:30:10:8:true LEADING:3 TRAILING:3 SLIDINGWINDOW:4:20 MINLEN:50). The processed reads with length greater than 50nt were defined as clean reads. Forward and reverse reads were C-to-T and G-to-A converted, respectively, and mapped to the appropriate converted reference using the meRanGh tool (align bsRNA-seq reads to reference using HiSat2) in MeRanTK (Rieder et al. 2016). Only uniquely mapped reads were retained and replaced by the original unconverted reads. Mapping parameters for each library are shown in Table S4.

#### Background removal, non-conversion site calling and site annotation

For transcriptome-wide candidate m^5^C site discovery, data from individual bsRNA-seq fraction libraries were pooled into their respective biological replicate (repB, repC, repE). To remove background non-conversion, reads containing more than three unconverted cytosines were removed from the bam files (‘**3C**’ filter; see also Table S4). Read counts at each cytosine position in the genome were obtained using the ‘mpileup’ function in samtools. Non-conversion sites were determined using a custom script with parameters ‘-minBQ 30 -- overhang 6’. We observed a cytosine bias of uncertain origin at the 5’ and/or the 3’ end of the reads and thus opted to mask non-converted cytosines within the terminal 6nt of each read to avoid overestimation of non-conversion. Candidate sites with a signal-to-noise ratio ≤ 0.9 (3C/raw; ‘**S/N≥0.9**’) were further flagged and suppressed. To retain high-confidence non-conversion sites, the following criteria were applied: (1) total read coverage of ≥ 30 (‘**≥30RC**’), (2) non-converted C of ≥ 5 (‘**5C**’), (3) C+T coverage ≥ 80% (‘**80CT**’). An average non-conversion of ≥ 10% across the biological triplicates (‘**10MM**’) was also required. Candidate sites were annotated to the longest transcript according to the GENCODE v28 annotation (UCSC) using a custom script and mapped to six features: 5’UTR, CDS, 3’UTR, intronic, ncRNA_exonic and ncRNA_intronic. The RNA transcript type for each candidate site was extracted simultaneously. All candidate sites that could not be annotated were considered to be intergenic.

#### Read mapping and non-conversion site calling in tRNAs and rRNAs

For the tRNA analysis, reads were mapped to the pre-tRNA reference using the meRanT tool within meRanTK (Rieder et al. 2016) with parameter (-k 10). Mapped reads containing more than three unconverted cytosines were removed from the bam files (3C filter) and only reads mapping to the predicted mature tRNA regions were retained (called ‘processed’ reads). tRNA sites were called as described for transcriptome-wide sites using only processed tRNA reads (*e.g.* sites shown in Figure S4C,D). For rRNA analysis, reads were mapped to the ribosomal DNA complete repeat unit using meRanT tool within meRanTK. rRNA sites were called as described for transcriptome-wide site and using all reads mapped to the rRNA reference, excluding reads containing more than three unconverted cytosines, i.e. potentially including unprocessed precursors. All rRNA-related sites are reported in Figure S4B and Table S5A, while only sites in mature rRNA regions are shown in Figure S3C.

### Candidate site characteristics and metagene analyses

#### Metagene distribution analysis of candidate m^5^C sites

Only exonic protein-coding candidate sites were used for distribution analysis along mRNAs. The relative position of each candidate site in the corresponding transcript feature (5’UTR, CDS or 3’UTR) was identified. For metagene density plots, each transcript feature was assigned a value corresponding to average feature length fraction out of the over-all transcript length. The background C control was generated using all C positions within the interrogated transcript segment of genes with candidate sites. Spatial enrichment analyses were conducted using RNAModR (Evers et al. 2016) with all unmodified C within the analysed region serving as background control. RNAModR was run using the GENCODE v20 annotation (UCSC) to build the transcriptome (DOI: <u>10.18129/B9.bioc.BSgenome.Hsapiens.UCSC.hg38</u>).

#### Motif enrichment analysis

To acquire the sequence preference proximal to candidate sites, 21nt sequences centred around each candidate site were extracted from the genome with BEDTools (Quinlan and Hall 2010). All C positions from genes with candidate sites were used as background control. Sequence logo plots were generated with ggseqlogo (Wagih 2017).

#### Secondary structure analysis

Secondary structure analysis surrounding the candidate mRNA sites was done as described previously (Huang et al. 2019). Specifically, 25nt sequences upstream and downstream of each candidate site were extracted from the genome with BEDTools (Quinlan and Hall 2010) and folded with the RNAfold tool in the ViennaRNA Package 2.0 (Lorenz et al. 2011) using default parameters. The percentage of paired bases at each position was calculated from the folding results.

#### Gene ontology (GO) analysis

Gene Ontology (GO) analysis was performed using the *enrichGO* functionality within the clusterProfiler package (Yu et al. 2012). Genes with FKPM ≥1 in the bsRNA-seq dataset were used as background control gene set. Resulting enriched GO terms were restricted to Bonferroni-corrected p-values < 0.05.

#### Codon position bias of candidate sites

The codon position of each candidate site was calculated using various numpy functions in python from arrays of codon counts for each transcript. Arrays were obtained by counting all codons with a candidate site at the first, second or third position and comparison to all used codons. The enrichment of each codon was calculated using Fisher’s exact test.

#### Translation efficiency and mRNA stability analyses

For analysis of translation efficiency and mRNA half-life, publicly available HeLa datasets GSE49339 (Wang et al. 2014) and GSE102113 (Arango et al. 2018) were downloaded from NCBI. For analysis of translation efficiency during the somatic cell cycle (Asynchronous, S phase and M phase) the publicly available dataset GSE79664 (Park et al. 2016) was also obtained. To quantify RNA or ribosome-protected fragment (RPF) abundance, annotations with more than 60 mapped reads were selected and normalized using the Trimmed Mean of M values (TMM) method implemented in the edgeR Bioconductor package (Robinson et al. 2010). Translation efficiency was calculated by dividing TMM normalized RPF values to that of RNA.

Cumulative density of translation efficiency and mRNA half-life were generated using genes containing candidate sites versus all expressed genes in HeLa cells.

#### Clustering of candidate sites across bsRNA-seq fractions

Only candidate sites in protein-coding genes were taken forward to clustering analyses. Candidate sites with any coverage in 9 out of 12 bsRNA-seq fraction samples and an average coverage ≥10 across the biological triplicates in each bsRNA-seq fraction were considered (F1234 clustering). As some candidate sites do not have enough coverage in bsRNA-seq fraction 1 (average coverage <10 in F1), clustering was performed again considering only bsRNA-seq fractions 2-4 (F234; see Figure S2). Average non-conversion level for each candidate site per bsRNA-seq fraction was used as input data. Clustering was performed using the Mfuzz soft clustering method (Kumar and M 2007) with the fuzzifier and cluster number parameters set to *m* = 2 and *c* = 9, respectively.

#### Logistic regression of non-conversion change between bsRNA-seq fractions

Sequential pairwise non-conversion level comparison between bsRNA-seq fractions F1 and F2, or F2 and F3, or F3 and F4 was carried out. Only sites with average coverage ≥10 in both bsRNA-seq fractions analysed were considered. Information from each fraction is specified (the average number of methylated Cs and average number of unmethylated Cs at a given site), and a logistic regression test was applied to compare the proportion of methylated Cs across two fractions using the methylKit R package (Akalin et al. 2012). Sites with a q-value <0.05 and relative methylation difference ≥10% were defined as differentially methylated sites.

#### Statistical analysis

All bioinformatics-associated statistical analyses (unless stated otherwise) were performed in ‘R’. p<0.05 is considered as statistically significant. Significance of average non-conversion changes were assessed by unpaired, two-tailed Students’ *t*-test. Significance for candidate site distributions across mRNA regions within sampling dependent pools (Figures 5,S10) was assessed using binomial testing.

All significance levels are: ns - not significant, * p<0.05, ** p<0.01, *** p<0.001, **** p<0.0001.

## Data Access

All source code used during this study is available at https://github.com/zhanghena/ bsRNA-seq-m5C/. All raw and processed sequencing data generated in this study have been submitted to the NCBI Gene Expression Omnibus (GEO; https://www.ncbi.nlm.nih.gov/geo/) under accession number GSE140995.

## Acknowledgements

The authors would like to thank Maurits Evers for assistance with and expansion of RNAModR; Hardip Patel for general assistance with R as well as statistical analyses; Andrew Shafik, Natalie J Beveridge, Brian J Parker, Jiayu Wen, Aaron L Statham, David T Humphreys and Jeffrey E Squires for contributions to early work that led to this study.

## Authors’ contributions

T.P., U.S., T.S. developed the study; U.S., T.S., A.P., S.G. performed experiments; H.-N.Z., A.H. performed bioinformatics analyses; L.Y., S.J.C. provided conceptual input; U.S., T.P. wrote the manuscript.

## Supplementary Figures

**Supplementary Figure S1:**
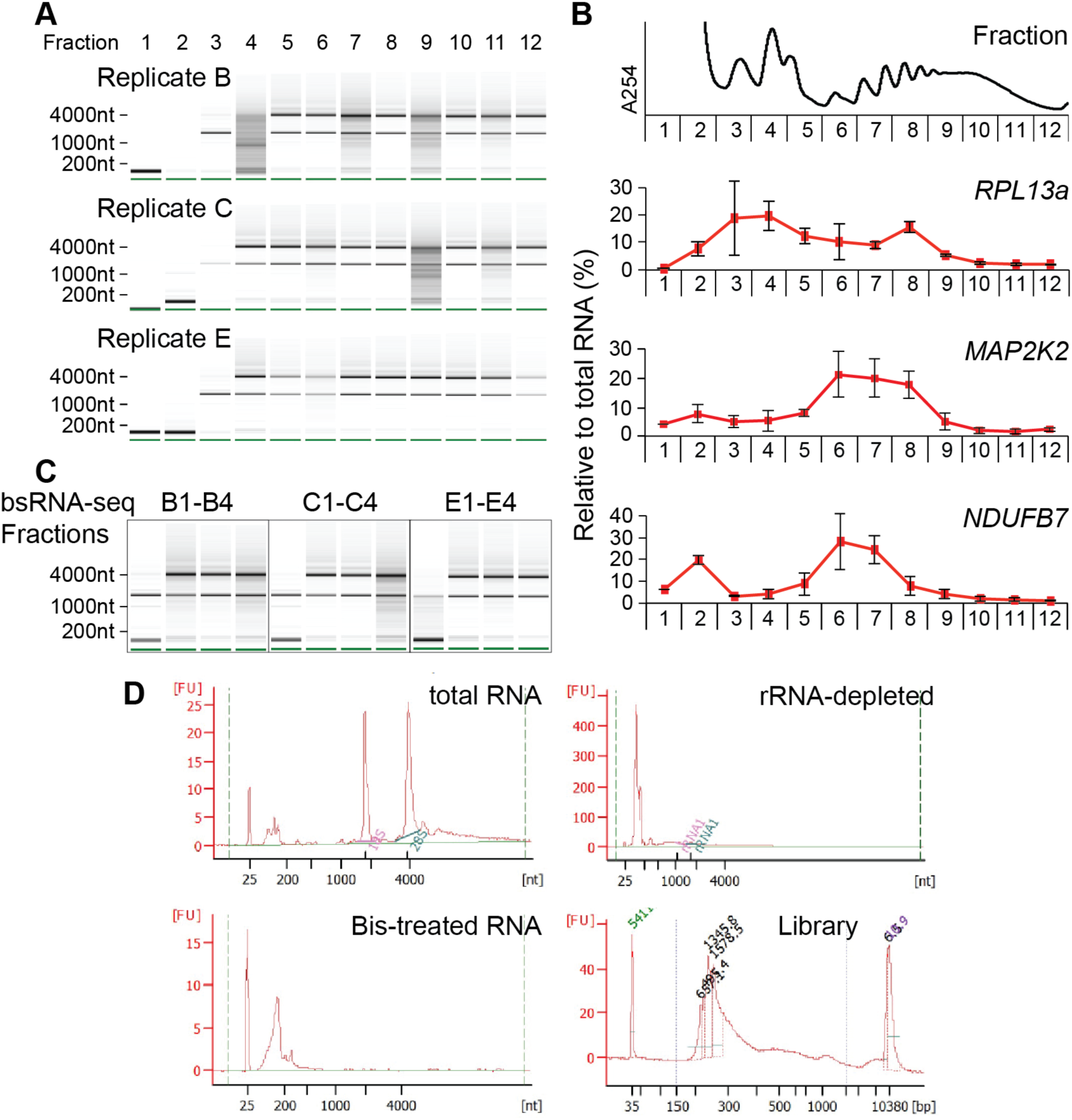
Quality controls for polysome profiling and bsRNA-seq sample preparation. Related to Figure 1. A: Distribution of tRNA and rRNA across gradients. Equal proportions of total RNA from each RNA fraction was analysed by microfluidic electrophoresis (Bioanalyzer RNA 6000 Nano Chip; equal proportions of recovered RNA were loaded). Pseudo-gel images for each of the three biological replicates are shown. B: Distribution of additional representative mRNAs across gradients. mRNA levels in each RNA fraction were determined by RT-qPCR. Results for three mRNAs of different coding region length are shown: *RPL13a* (ribosomal protein L13a), *MAP2K2* (mitogen-activated protein kinase kinase 2) and *NDUFB7* (NADH:ubiquinone oxidoreductase subunit B7). mRNA levels per fraction were normalised to the level of a spike-in control, rescaled as percentage of total signal across all fractions, and are shown as mean ± standard deviation across the three biological replicates. A representative absorbance trace (254nm) is shown at the top for reference. C: RNA quality of bsRNA-seq fractions prior to bisulfite treatment. RNA from each bsRNA-seq fraction was analysed by microfluidic electrophoresis (Bioanalyzer RNA 6000 Nano Chip; an equal amount of RNA was loaded per well). Pseudo-gel images for each of the three biological replicates are shown. D: Microfluidic electrophograms for biological replicate E tracing the RNA quality at each step from input to the final library (from left to right). Data shown are exemplary for all biological replicates.

**Supplementary Figure S2:**
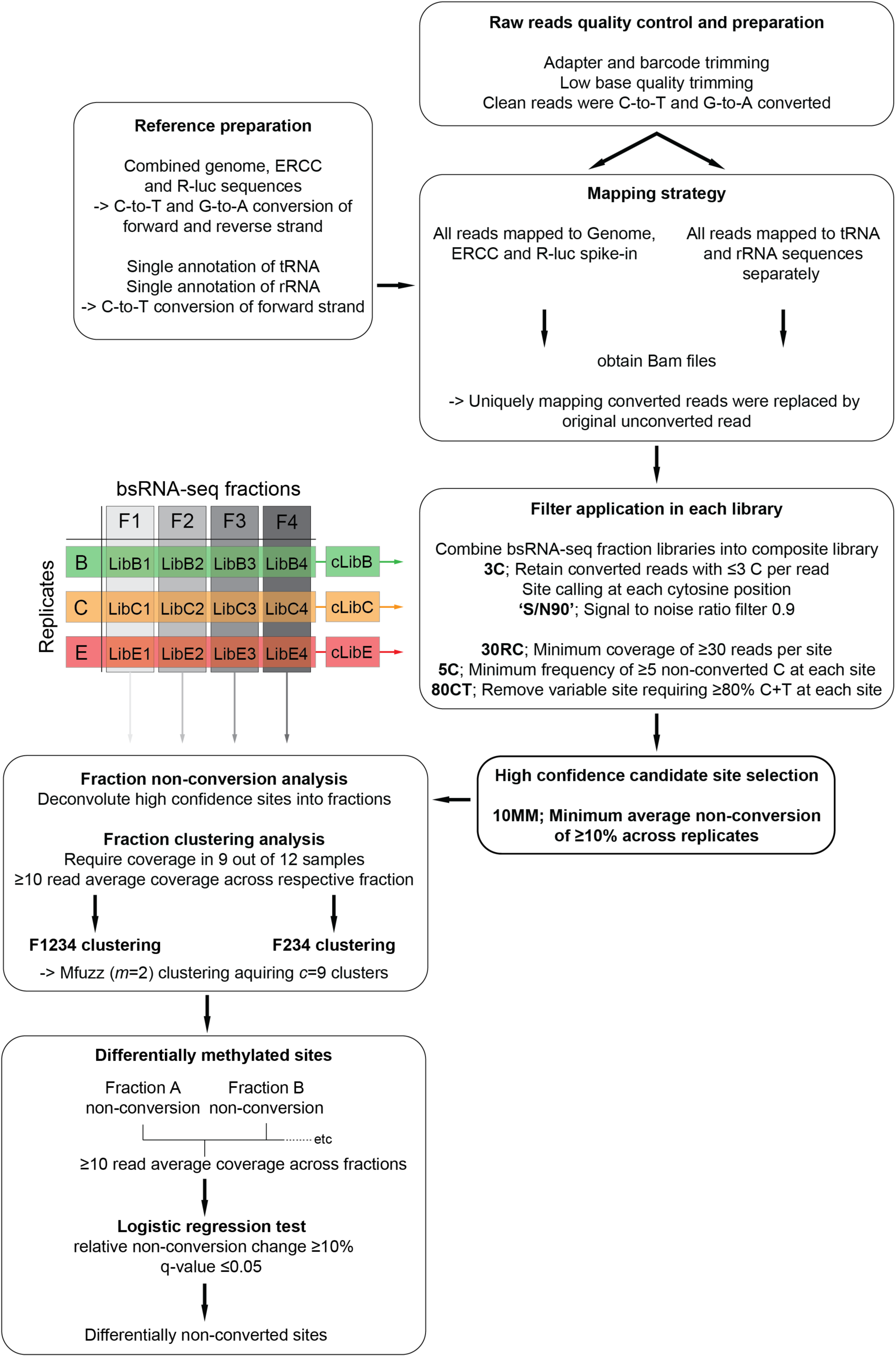
bsRNA-seq mapping and data analysis. Related to Figures 2 and 5. Workflow from bsRNA-seq read processing and mapping, m^5^C candidates site selection to clustering by non-conversion level across polysome gradients. For the definitive site selection, steps in the workflow were performed sequentially. Selection criteria for high confidence candidate sites and alternate groupings of bsRNA-seq libraries for different purposes are indicated. Note, four bsRNA-seq fraction libraries representing distinct translation states were sequenced per biological replicate, creating a total of twelve libraries termed **LibB1-4**, **LibC1-4** and **LibE1-4**. For global m^5^C candidate site calling, Libs 1-4 were combined into one composite library for each biological replicate, creating **cLibB**, **C** and **E**. These composite libraries approximate a total transcriptome-wide survey for each biological replicate. For clustering analyses, libraries from corresponding bsRNA-seq fractions (i.e. **LibB1**, **LibC1** and **LibE1** and so forth) formed biological replicates of each other.

**Supplementary Figure S3:**
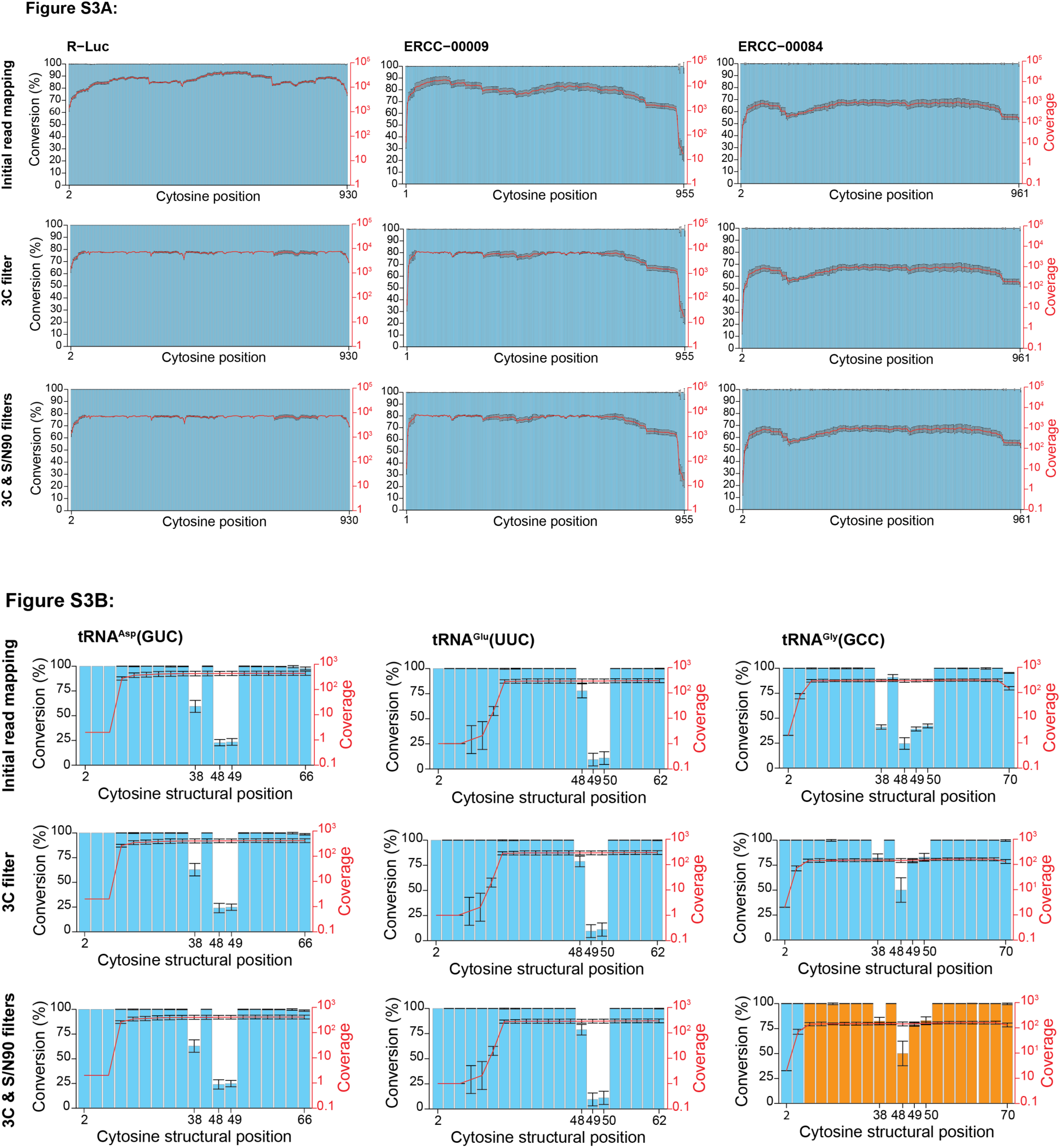
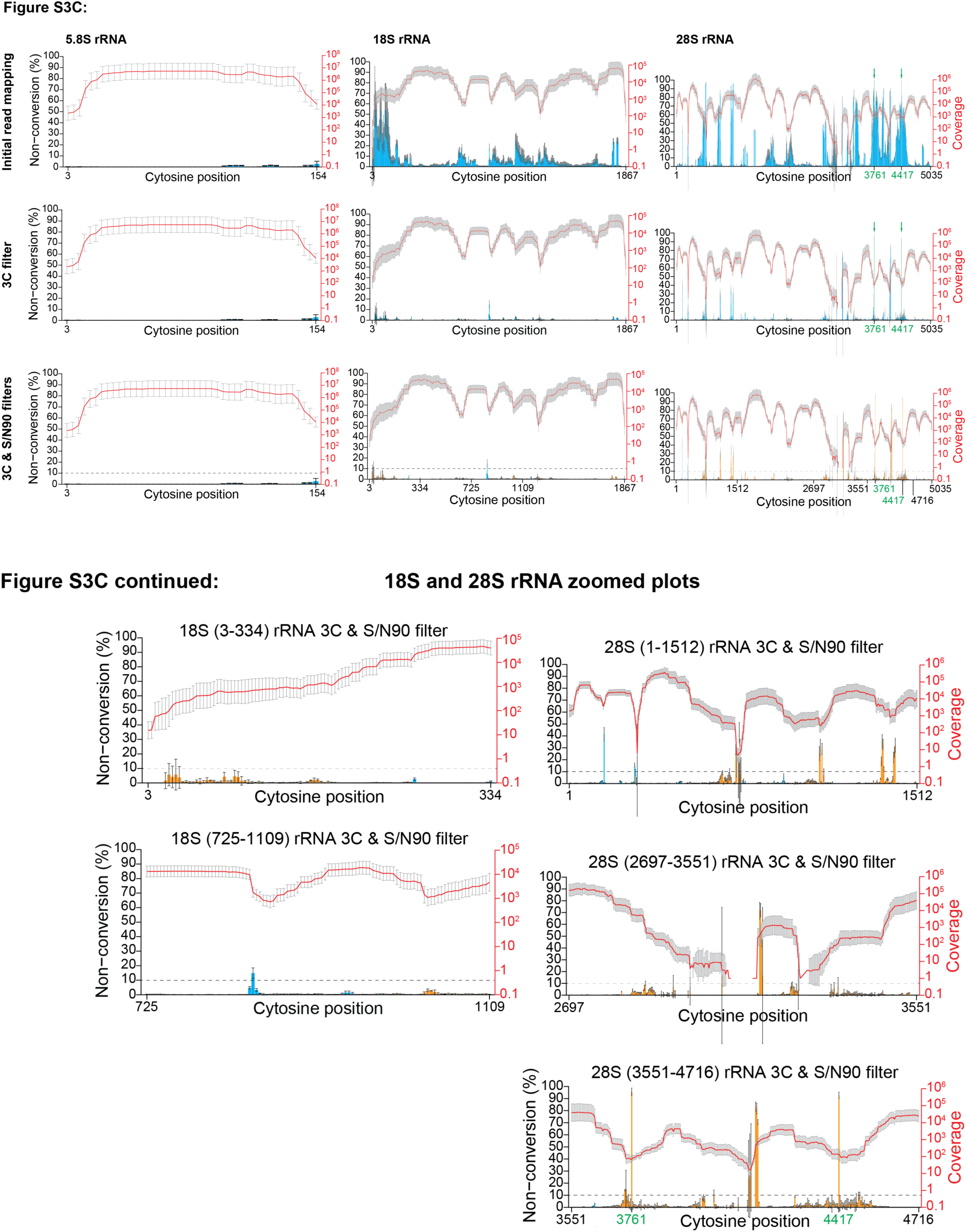
Effects of the 3C and S/N90 filters on specificity and sensitivity of m^5^C candidate site detection. Related to Figure 2. In each panel, plots are arranged vertically by RNA under investigation, and horizontally by the extent of sequential filtering (initial read mapping—after removing reads with >3 non-converted cytosines ‘3C filter’—after suppressing sites below the chosen signal-to-noise threshold ‘3C & S/N90 filter’ [less than 90% of reads passing the 3C filter]). Dual y-axis plots show either cytosine conversion (A,C) or non-conversion (B) (left y-axis, blue bars) and read coverage (right y-axis, red line) against cytosine position in the respective reference sequence (x-axis). Data is shown as mean across the three biological replicates with error bars indicating ± standard deviation. Candidate sites disqualified by the S/N90 filter are identified by orange bars. The effects of the filters were evaluated using selected spike-in control (A), rRNA (B) and tRNA (C) sequences. A: Panel of spike-in controls, *R-Luc* RNA and two arbitrarily selected ERCC transcripts. B: Mature ribosomal RNA species. Note that cytosine non-conversion is plotted for improved visualisation. The fourth to sixth panels show zoomed-in plots of fully filtered 18S and 28S rRNA data. Residues of zoomed regions are indicated on the top and correspond to numbering in full-scale plots. Green arrows and position labelling indicate the two known m^5^C sites in 28S rRNA (Sharma et al. 2013). C: Selected tRNA examples. tRNA^Asp^(GUC), tRNA^Glu^(UUC) and tRNA^Gly^(GCC) were chosen to represent different m^5^C positions within tRNAs and to illustrate the adverse effect of the chosen filters on tRNAs with >3 modified cytosines. Cytosine numbering is according to the tRNA consensus structural positions.

**Supplementary Figure S4:**
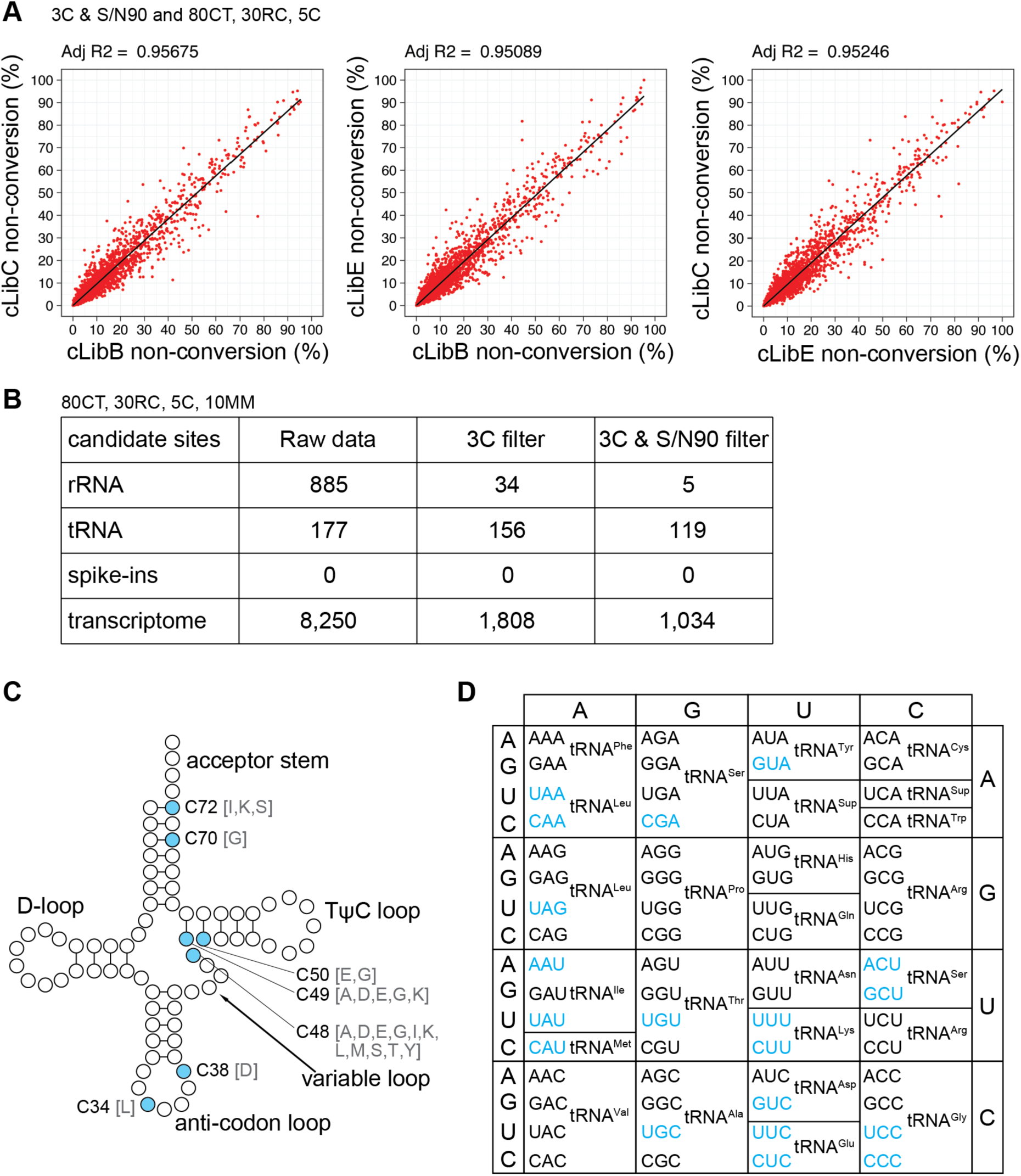
m^5^C candidate site call reproducibility across biological replicates and effects of non-conversion ‘noise suppression’. Related to Figure 2. A: Pair-wise scatter plot comparisons of transcriptome-wide candidate sites called in composite libraries of each biological replicate. Sites shown passed the 80CT, 30RC and 5C filter in their respective composite library (a non-conversion cut-off was not applied). Further to that, only sites with coverage in all three replicates were used. The adjusted R-squared value following linear regression is shown. B: Effect of the 3C and S/N90 filters on candidate site calling in different RNA types. The number of candidate sites that passed the 80CT, 30RC, 5C filter in their respective composite library and fulfilled the 10MM criterion are listed. C: Position of candidate sites in the tRNA cloverleaf consensus structure. Each circle indicates a nucleotide position within the tRNA cloverleaf structure, with blue filled circles indicating position at which candidate sites were identified. Iso-acceptors found to carry the candidate site are identified by the single letter amino acid code. D: Genetic code table highlighting tRNA iso-decoders with candidate sites in blue. C-D: Of note, we detected the NSUN2-dependent sites at the edge of the variable loop at position C48-50 in a variety of tRNA iso-decoders, as well as at position C34 of intron-containing tRNA^Leu^(CAA). We further identified the TRDMT1-dependent modification of C38 in tRNA^Asp^(GUC). Interestingly, we also detected several candidate sites at structural position C72. The established NSUN6-dependent sites in tRNA^Thr^(UGU) and tRNA^Cys^(GCA) iso-decoders did not receive read coverage. Instead, we saw clear non-conversion at C72 in tRNA^Ile^(UAU), tRNA^Lys^(CUU) and tRNA^Ser^(ACU); these might be novel NSUN6 substrates. C70 in tRNA^Gly^(CCC) is indicated in the figure, however, detection of this site is heavily driven by the terminal base of reads in one direction, thus likely suspect.

**Supplementary Figure S5:**
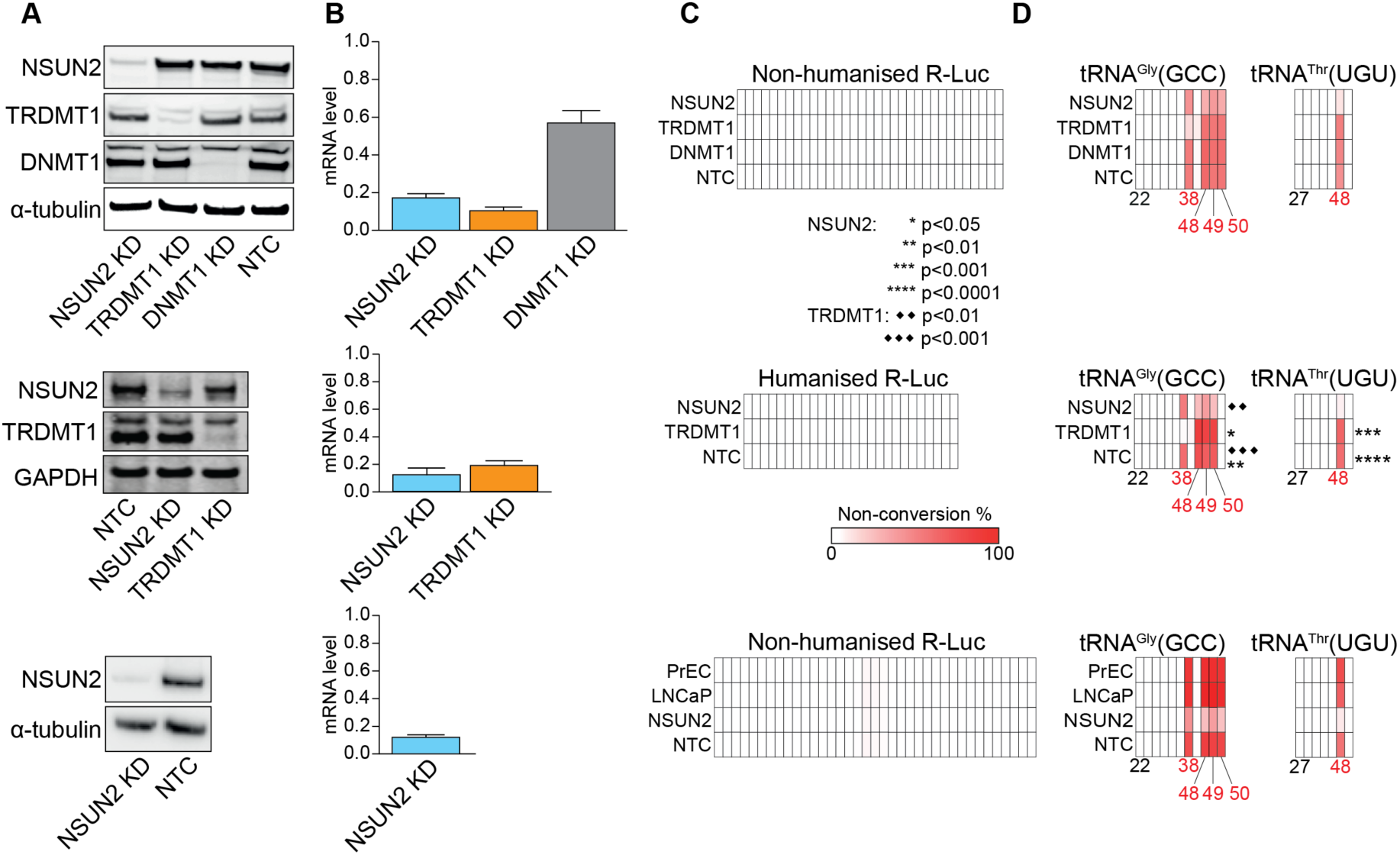
siRNA knockdown efficiency controls. Related to Figure 3. HeLa cells, or the prostate cell lines LNCaP or PrEC, were transiently transfected with siRNAs targeting the m^5^C:RNA methyltransferases NSUN2 or TRDMT1, the m^5^C:DNA DNMT1 or a non-targeting control (NTC) as indicated in the panels. Lysates were prepared for western analysis (panel A). Total RNA was extracted for RT-qPCR (panel B) and amplicon-bsRNA-seq (panel C,D), respectively. Across panels, results are arranged with HeLa ‘confirmatory’ data on top (N=1), HeLa ‘in depth’ data in the middle (N=3), and prostate cell line ‘confirmatory’ data at the bottom (N=1). A: Methyltransferase and control protein levels after siRNA knockdown. Western blots for NSUN2, TRDMT1, DNMT1 and the internal controls alpha-tubulin or GAPDH are indicated on the left. siRNA knockdown condition is shown below. One replicate for the HeLa ‘in depth’ data is shown; similar results were obtained for the other replicates. B: Methyltransferase mRNA levels after specific siRNA knockdown are shown relative to those in the NTC control. HeLa RT-qPCR data were normalised to the internal control genes *HPRT* (top) or *GAPDH* (middle), RT-qPCR data from the prostate cell lines were normalised to the geometric mean of the internal control genes *MRPL9*, *H2AFV* and *TCF25*. ‘Confirmatory’ data (top and bottom) are shown as averages of three technical replicates. ‘In depth’ data (middle) are shown as averages of three biological replicates. Error bars indicate standard error of the mean. C-D: Associated amplicon-bsRNA-seq results for the *R-Luc* spike-in negative controls (C) and selected tRNAs as positive controls (D). The interrogated RNA is indicated above each grid. Grids are organised by knockdown sample (in rows; siRNA target indicated on the left) and cytosine position along the analysed transcript section (in columns; for tRNAs structural position coordinates are given for the first interrogated cytosine [in black], as well as the candidate m^5^C sites [in red]). A white-to-red colour scale (shown in the middle row) is used to tint each square by the degree of cytosine non-conversion. Note, C38 in tRNA^Gly^(GCC) is a known target of TRDMT1, whereas C48-50 in tRNA^Gly^(GCC) and C48 in tRNA^Thr^(UGU) are mediated by NSUN2. Student’s *t*-test results of non-conversion change for tRNA^Gly^(GCC) in the ‘confirmatory’ HeLa data is indicated next to the grid with asterisks and diamonds showing significance to the NSUN2 or TRDMT1 knockdown sample, respectively (key in top panel).

**Supplementary Figure S6:**
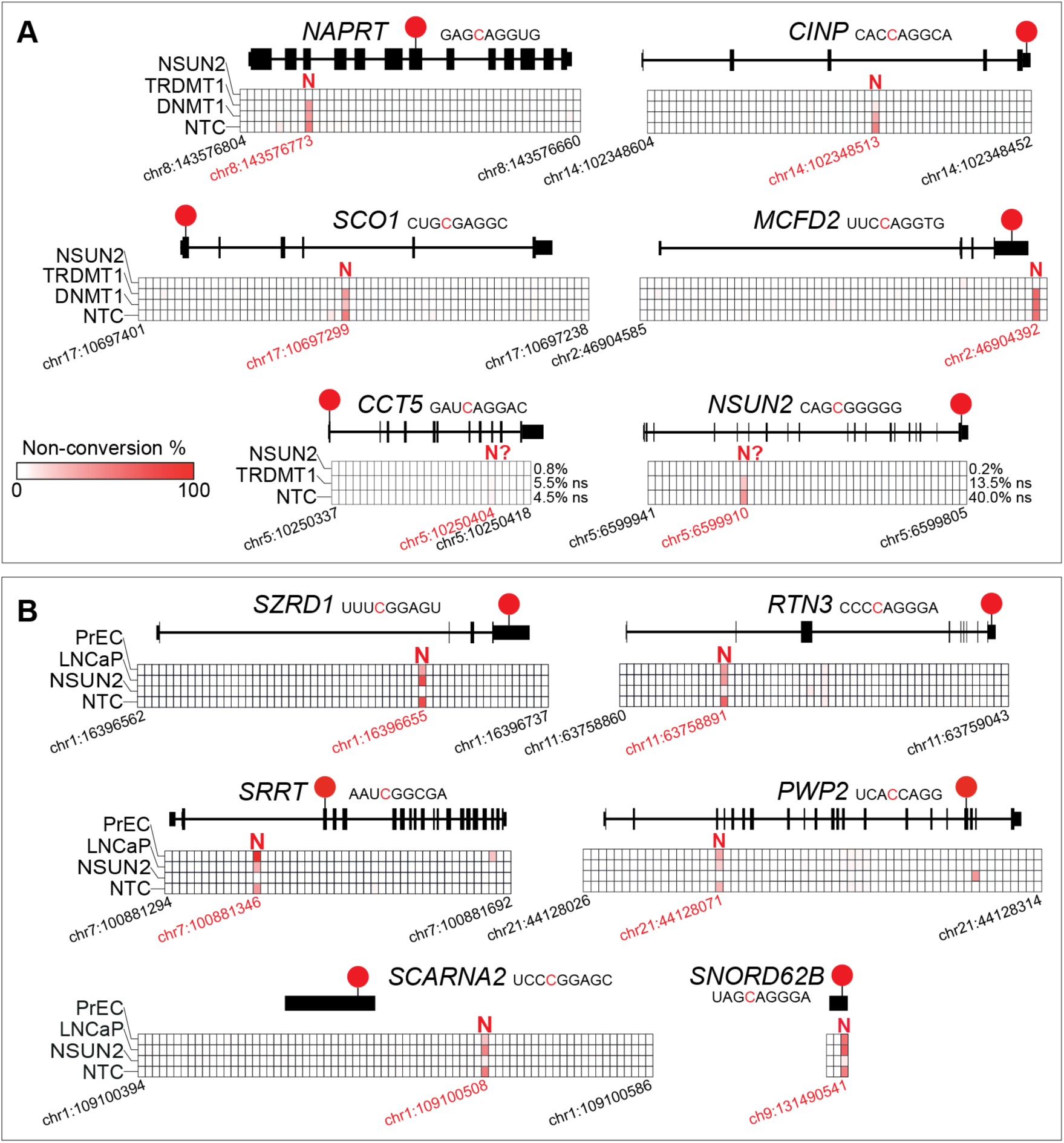
Additional validation and NSUN2-dependence of candidate m^5^C sites in mRNA and ncRNA. Related to Figure 3. Amplicon-specific bsRNA-seq was performed with total RNA isolated from HeLa cells, or the prostate cell lines LNCaP or PrEC, after siRNA-mediated m^5^C:RNA methyltransferase knockdown targeting NSUN2 or TRDMT1 along with control siRNAs (targeting m^5^C:DNA Methyltransferase DNMT1 or a non-targeting control [NTC]; see Figure S5 for knockdown efficiency controls). Read coverage achieved per amplicon was from 3,500 to 67,500 (see Table S6 for details). Grids in the centre of each diagram are organised by knockdown sample (in rows; siRNA target indicated on the left) and cytosine position along the analysed transcript section (in columns; genomic coordinates are given for the first and last interrogated cytosine position [in black], as well as the candidate m^5^C site [in red]). A white-to-red colour scale (shown in panel A) is used to tint each square by the degree of cytosine non-conversion. The longest isoform of the interrogated mRNA (based on Ensembl) is shown as a pictogram above the grid. Candidate m^5^C site position is indicated by a red circle and its sequence context (non-converted cytosine indicated in red) given alongside the gene name. The enzyme identified to be responsible for cytosine methylation is indicated above the candidate sites: N – NSUN2; N? – unresolved but likely NSUN2. A: ‘Confirmatory’ HeLa cell data (N=1) is shown for mRNA sites in *NAPRT* (nicotinate phosphoribosyltransferase), *CINP* (cyclin dependent kinase 2 interacting protein), *SCO1* (SCO cytochrome C oxidase assembly protein 1) and *MCFD2* (multiple coagulation factor deficiency 2) (top and middle row). ‘In depth’ data (N=2-3) is shown as the average of at least two biological replicates for mRNA sites in *CCT5* (chaperonin containing TCP1 Subunit 5) and *NSUN2* (bottom row). These two sites are likely controlled by NSUN2, although the non-conversion change is not significantly (ns) different as determined by Student’s *t*-test on the means in comparison to the NSUN2 KD sample (Obtained non-conversion averages are indicated to the right of the grid; see also Table S6). B: ‘Confirmatory’ prostate cell line data (N=1) is shown for mRNA sites in *SZRD1* (SUZ RNA binding domain containing 1), *RTN3* (reticulon 3), *PWP2* (PWP2 small subunit processome component) and *SRRT* (serrate) (top and middle row) and ncRNA sites in *SCARNA2* (small Cajal body-specific RNA 2) and *SNORD62B* (small nucleolar RNA, C/D box 62B) (bottom row).

**Supplementary Figure S7:**
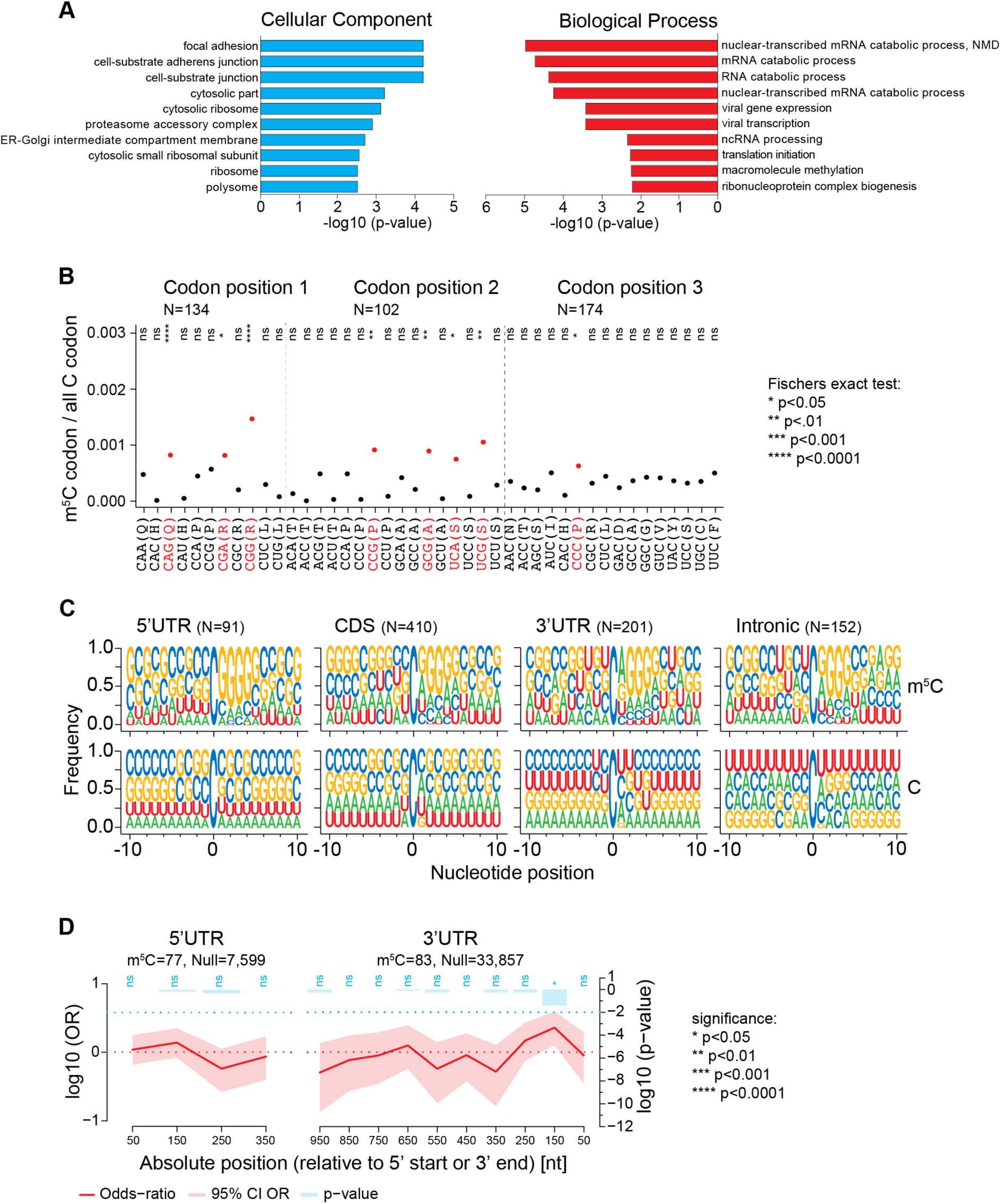
Sequence context and NSUN2-dependence of candidate m^5^C sites. Related to Figure 4. A: Gene Ontology (GO) term enrichment of candidate sites. Analysis was performed using the enrichGO function in ClusterProfiler using transcriptome-wide candidate sites (n=846), with all genes detected at FKPM≥1 in the bsRNA-seq used as background. Bonferroni correction was applied and terms with p<0.05 deemed enriched. The ten most enriched terms are shown (for full list of enriched terms see Table S7). Enrichment for Cellular Component (left) and Biological Process (right) are shown, no enrichment was obtained for Molecular Function GO terms. B: Codon position enrichment analysis of candidate sites with the CDS of protein-coding genes. All three codon positions were analysed and are indicated at the top. Codons preferentially containing candidate sites are indicated in red, with significance following Fisher’s exact test indicated: ns - not significant, * <0.05, ** <0.01, *** <0.001. C: Sequence context of candidate sites (top) in comparison to all cytosines in the same transcripts (bottom). Logos were generated for candidate sites present within the four RNA transcript regions (5’UTR: m^5^C, N=91; Null=150,943. CDS: m^5^C, N=410; Null=1,139,649. 3’UTR: m^5^C, N=201; Null=589,922. Intronic: m^5^C, N=152; Null=26,326,026). All sites from the ‘protein-coding’ (N=846) and ‘NMD’ RNA biotype (N=8) were included. D: Spatial enrichment analyses of candidate sites within mRNAs. Site are placed into bins as indicated on the x-axis. Site distribution across bins is compared to matching randomised cytosine sampling (Null) and the log10 Odds-ratio (OR) is plotted as a red line with the 95% confidence interval (CI) shaded. Significance of enrichment is plotted as the log10 p-value by blue bars: ns - not significant, * <0.05, ** <0.01, *** <0.001. Analyses were anchored at either the transcription start (left) or the transcription end (right) and performed using RNAModR with a bin width of 100nt and a window of 400nt for the 5’UTR or 1000nt for the 3’UTR region.

**Supplementary Figure S8:**
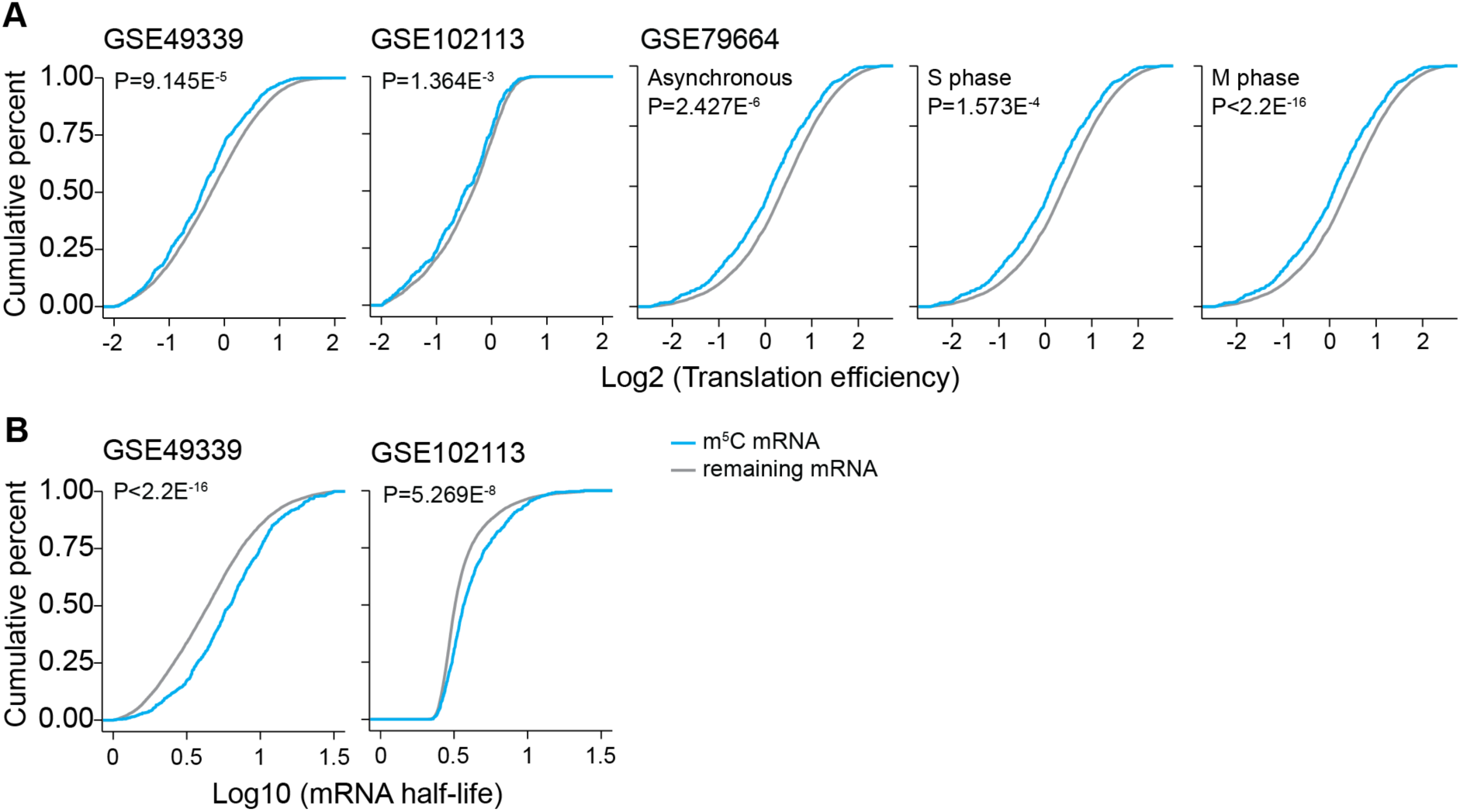
Correlation of mRNA translation and stability with m^5^C site content. Related to Figure 4. A: Cumulative density distribution of translation efficiency for candidate site-containing and all remaining mRNAs. Translation efficiency values are based on HeLa cell ribosome profiling data from [Wang et al. (2014); first plot], [(Arango et al. 2018); second plot] or [(Park et al. 2016); cell cycle plots]. (GSE49339: m^5^C mRNA, N=666; remaining mRNA, N=12,185. GSE102113: m^5^C mRNA, N=667; remaining mRNA, N=11,460. GSE79664: m^5^C mRNA, N=669; remaining mRNA, N=11,717) B: Cumulative density distribution of mRNA half-life for candidate site-containing and all remaining mRNAs. HeLa cell mRNA half-life data was taken from [(Wang et al. 2014); first plot)] or [(Arango et al. 2018); second plot]. (GSE49339: m^5^C mRNA, N=649; remaining mRNA, N=10,569. GSE102113: m^5^C mRNA, N=612; remaining, N=11,175).

**Supplementary Figure S9:**
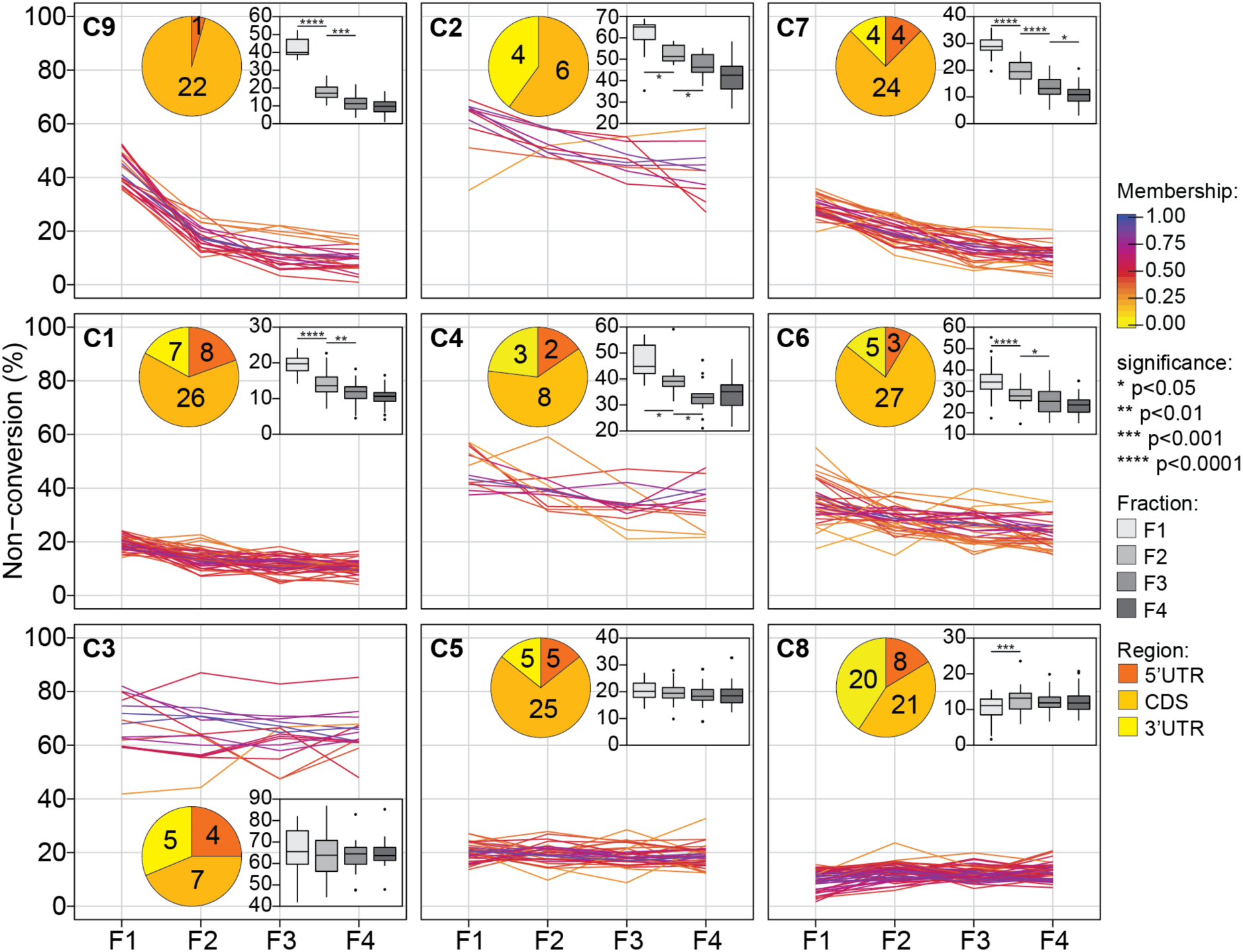
Clustering of candidate m^5^C site non-conversion level patterns across the polysome profile. Related to Figure 5. bsRNA-seq libraries were grouped as biological triplicates per fraction (e.g. LibB 1, LibC1 and LibE1 each report on sites detected in bsRNA-seq fraction 1), allowing the calculation of average non-conversion levels per individual site and per fraction. Non-conversion levels per individual site across the polysome profile were partitioned into nine soft clusters (C1-9 arranged here by similarity in trend with polysome profile) using Mfuzz. 254 candidate m^5^C sites were included based on having coverage in at least 9 out of 12 bsRNA-seq fraction samples and ≥10 average coverage in each of the four bsRNA-seq fractions. Degree of cluster ‘membership’ is indicated by the colour scale depicted to the right of panel A. Candidate sites with high membership (blue) have the best match to the respective cluster’s overall pattern. The legend to the right gives a colour/significance key applicable to all panels. Insets are: pie charts showing distribution of cluster members across different mRNA regions; boxplots showing site non-conversion distribution of cluster members across the polysome profile. Asterisks indicate significance p-value from unpaired, two-tailed Student’s *t*-test comparing the means of adjacent fractions. For Figure 5, clusters C9,2,7,1,4,6 were combined into the negative trend category, clusters C3,5 into the neutral category, and cluster C8 formed the positive category.

**Supplementary Figure S10:**
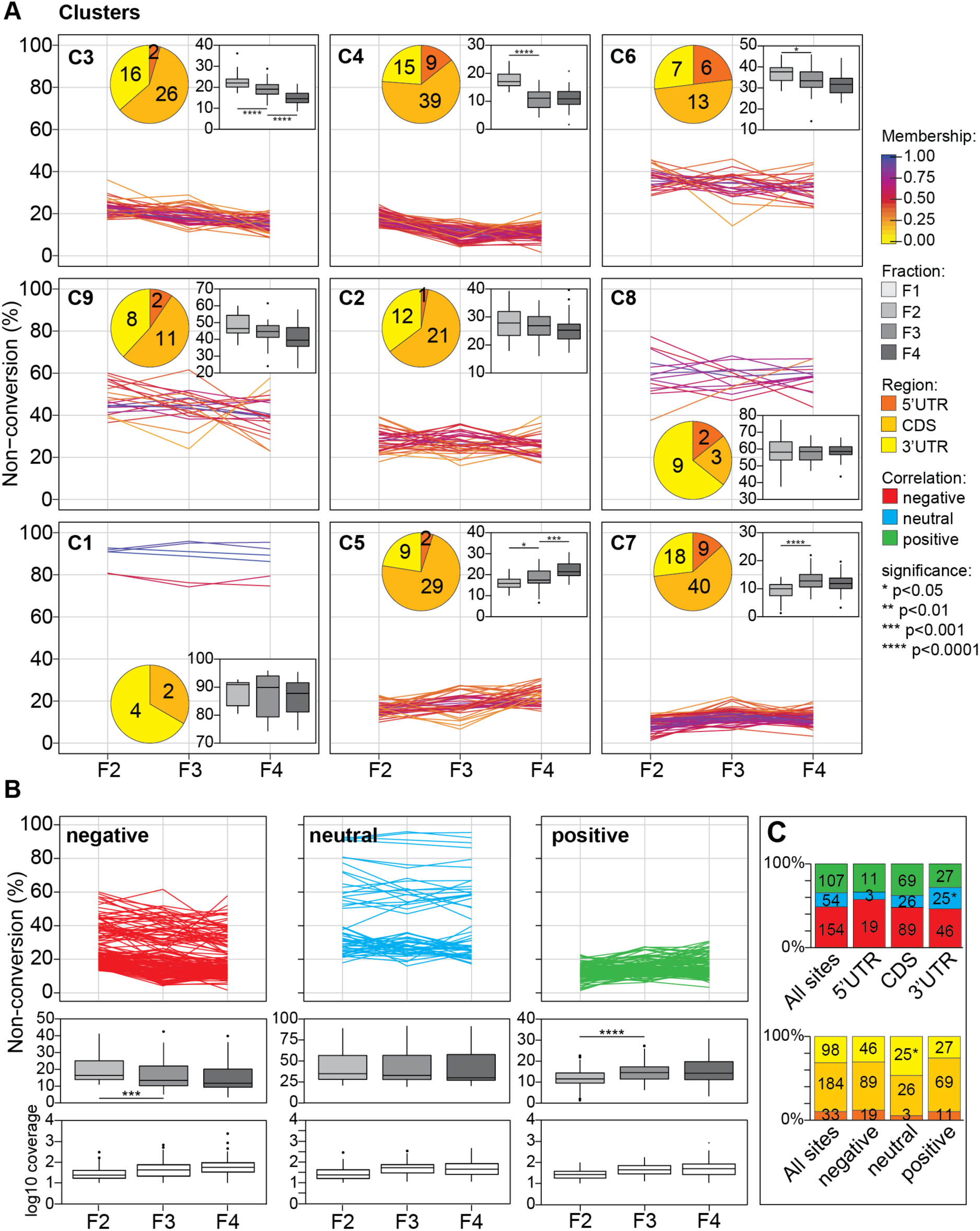
Relationship between non-conversion level and mRNA translation state for candidate m^5^C sites with insufficient coverage in bsRNA-seq fraction 1. Related to Figure 5. bsRNA-seq libraries were grouped as biological triplicates per fraction (e.g. LibB 1, LibC1 and LibE1 each report on sites detected in bsRNA-seq fraction 1), allowing the calculation of average non-conversion levels per individual site and per fraction. A: Non-conversion levels per individual site across the polysome profile were partitioned into nine soft clusters (C1-9 arranged here by similarity in trend with polysome profile) using Mfuzz. 315 candidate m^5^C sites were included based on having coverage in at least 9 out of 12 bsRNA-seq fraction samples and ≥10 average coverage in bsRNA-seq fractions 2-4 but failing this criterion for fraction 1. Degree of cluster ‘membership’ is indicated by the colour scale depicted to the right of panel A. Candidate sites with high membership (blue) have the best match to the respective cluster’s overall pattern. The legend to the right of panel A gives a colour/significance key applicable to all panels. Insets are: pie charts showing distribution of cluster members across different mRNA regions; boxplots showing site non-conversion distribution of cluster members across the polysome profile. Asterisks indicate significance p-value from unpaired, two-tailed Student’s *t*-test comparing the means of adjacent fractions. B: Mfuzz clusters from panel A were grouped into three translation state trend categories by visual inspection, showing a negative (clusters C3,4,6,9; N=133), neutral (clusters C2,8,1; N=75) or positive trend (clusters C5,7; N=107) with polysome association. Top panels: line graphs displaying individual site average non-conversion levels across fractions. Middle panels: boxplots showing distribution of site non-conversion levels in each fraction. Asterisks indicate significance p-value from unpaired, two-tailed Student’s *t*-test comparing the means of adjacent fractions. Bottom panels: boxplots showing distribution of site coverage in each fraction. C: Stacked bar charts showing distribution of sites in the different translation state trend categories across mRNA regions (top) and distribution of sites in different mRNA regions across translation state trend categories (bottom). Asterisks indicate significance p-values following binomial test against the distribution of all sites. The legend to the right of panel A gives a colour/significance key applicable to all panels.

**Supplementary Figure S11:**
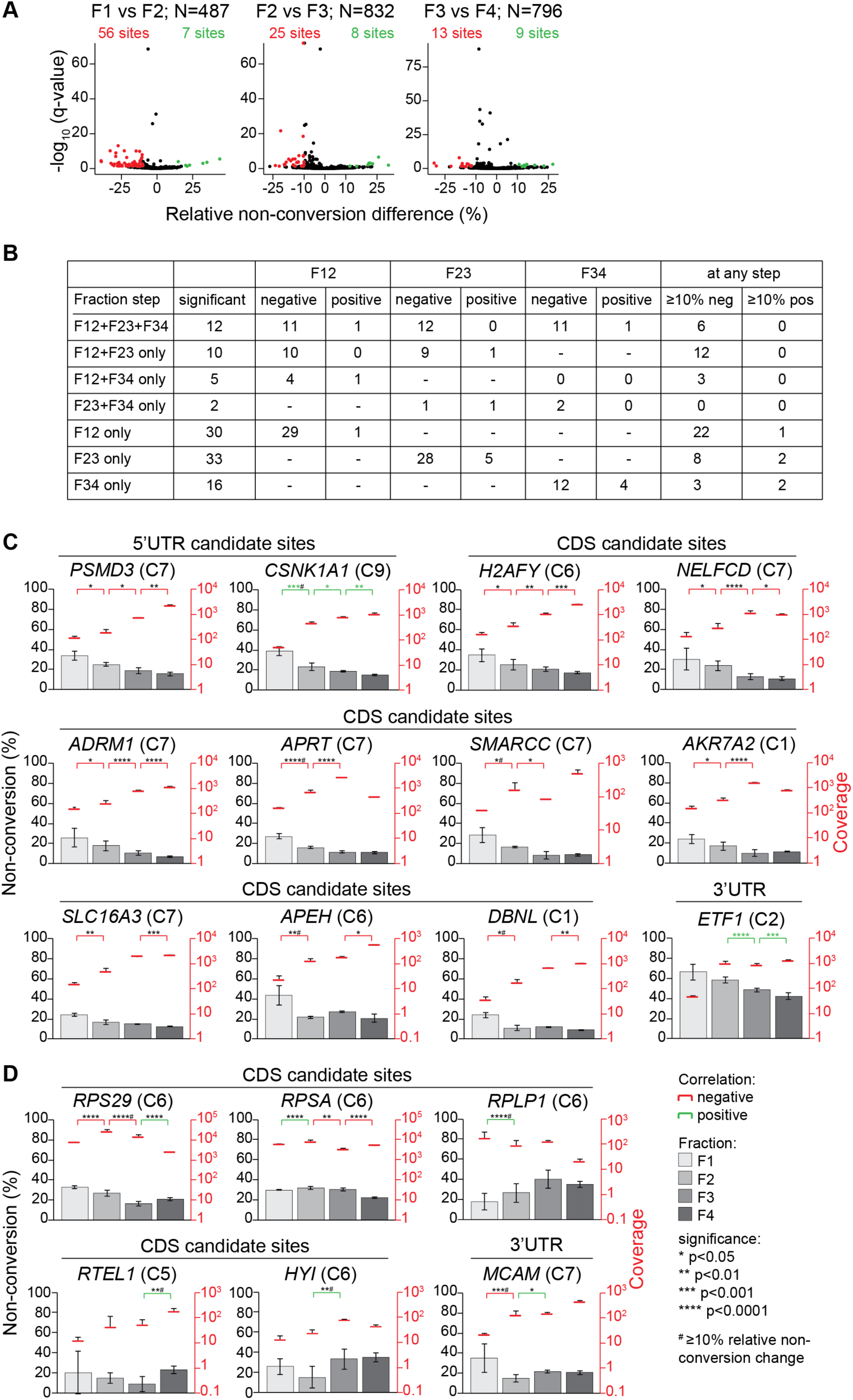
Identification of individual sites with significant correlation of cytosine non-conversion with polysome co-sedimentation. Related to Figure 6. A: Transcriptome-wide candidate m^5^C sites were used as input for logistic regression analysis (methylKit; (Akalin et al. 2012). For each pairwise comparison of adjacent polysome profile fractions a minimum of ≥10 average read coverage across each bsRNA-seq fraction was required. Plots show the -log10 q-value (FDR) of the non-conversion change, against the relative non-conversion difference for each pairwise comparison, with the number of sites that qualified indicated above (993 of 1,034 sites were involved in at least one such comparison). Each dot represents a single site, sites that score as significant (q-value <0.05) and showing a relative non-conversion change ≥10% are coloured in red (negative correlation) and green (positive correlation), respectively. B: Characteristics of sites included in the F1234 clustering (Figures 5,S9) that showed any significant trend as analysed in panel A. C-D: Examples of individual sites in mRNA showing significant correlation of cytosine non-conversion with translation state. Dual axis charts show cytosine non-conversion (bars) and coverage (red lines) for a given site across the polysome gradient. Data is shown as means across biological triplicates, with error bars indicating standard deviation. Asterisks indicate significance p-values after logistic regression testing (see key next to panel D). The gene name for each candidate site and its position within the mRNA is given. C: Individual examples with significant negative non-conversion change in at least two fraction steps. The cluster number from the F1234 clustering is indicated in brackets. Candidate sites shown here are from the following mRNAs: *PSMD3* (proteasome 26S subunit); *CSNK1A1* (casein kinase 1 alpha 1); *H2AFY* (H2A histone family member Y); *NELFCD* (negative elongation factor complex member C/D); *ADRM1* (adhesion regulating molecule 1); *APRT* (adenine phosphoribosyltransferase); *SMARCC* (SWI/SNF related, matrix associated, actin dependent regulator of chromatin subfamily C member 1); *AKR7A2* (aldo-keto reductase family 7 member A2); *SLC16A3* (solute carrier family 16 member 3); *APEH* (acylaminoacyl-peptide hydrolase); *DBNL* (drebrin like); *ETF1* (eukaryotic translation termination factor 1). D: Examples with significant positive non-conversion change in at least one fraction step. Candidate sites shown here are from the following mRNAs: *RPS29* (ribosomal protein 29); *RPSA* (ribosomal protein SA); *RPLP1* (ribosomal protein lateral stalk subunit P1); *RTEL1* (regulator of telomerase elongation helicase 1); *HYI* (hydroxypyruvate isomerase); *MCAM* (melanoma cell adhesion molecule).

## References

Aguilo F, Li S, Balasubramaniyan N, Sancho A, Benko S, Zhang F, Vashisht A, Rengasamy M, Andino B, Chen CH et al. 2016. Deposition of 5-Methylcytosine on Enhancer RNAs Enables the Coactivator Function of PGC-1alpha. Cell Rep 14: 479–492.

Akalin A, Kormaksson M, Li S, Garrett-Bakelman FE, Figueroa ME, Melnick A, Mason CE. 2012. methylKit: a comprehensive R package for the analysis of genome-wide DNA methylation profiles. Genome Biol 13: R87.

Amort T, Rieder D, Wille A, Khokhlova-Cubberley D, Riml C, Trixl L, Jia XY, Micura R, Lusser A. 2017. Distinct 5-methylcytosine profiles in poly(A) RNA from mouse embryonic stem cells and brain. Genome Biol 18: 1.

Anderson P, Ivanov P. 2014. tRNA fragments in human health and disease. FEBS Lett 588: 4297–4304.

Arango D, Sturgill D, Alhusaini N, Dillman AA, Sweet TJ, Hanson G, Hosogane M, Sinclair WR, Nanan KK, Mandler MD et al. 2018. Acetylation of Cytidine in mRNA Promotes Translation Efficiency. Cell 175: 1872–1886 e1824.

Baker SC, Bauer SR, Beyer RP, Brenton JD, Bromley B, Burrill J, Causton H, Conley MP, Elespuru R, Fero M et al. 2005. The External RNA Controls Consortium: a progress report. Nat Methods 2: 731–734.

Begik O, Lucas MC, Liu H, Ramirez JM, Mattick JS, Novoa EM. 2019. Integrative analyses of the RNA modification machinery reveal tissue- and cancer-specific signatures. bioRxiv doi:10.1101/830968: 830968.

Bhattacharyya SN, Habermacher R, Martine U, Closs EI, Filipowicz W. 2006. Relief of microRNA-mediated translational repression in human cells subjected to stress. Cell 125: 1111–1124.

Bian K, Lenz SAP, Tang Q, Chen F, Qi R, Jost M, Drennan CL, Essigmann JM, Wetmore SD, Li D. 2019. DNA repair enzymes ALKBH2, ALKBH3, and AlkB oxidize 5-methylcytosine to 5-hydroxymethylcytosine, 5-formylcytosine and 5-carboxylcytosine in vitro. Nucleic Acids Res 47: 5522–5529.

Blanco S, Bandiera R, Popis M, Hussain S, Lombard P, Aleksic J, Sajini A, Tanna H, Cortes-Garrido R, Gkatza N et al. 2016. Stem cell function and stress response are controlled by protein synthesis. Nature 534: 335–340.

Blanco S, Dietmann S, Flores JV, Hussain S, Kutter C, Humphreys P, Lukk M, Lombard P, Treps L, Popis M et al. 2014. Aberrant methylation of tRNAs links cellular stress to neuro-developmental disorders. EMBO J 33: 2020–2039.

Blanco S, Frye M. 2014. Role of RNA methyltransferases in tissue renewal and pathology. Curr Opin Cell Biol 31: 1–7.

Boccaletto P, Machnicka MA, Purta E, Piatkowski P, Baginski B, Wirecki TK, de Crecy-Lagard V, Ross R, Limbach PA, Kotter A et al. 2018. MODOMICS: a database of RNA modification pathways. 2017 update. Nucleic Acids Res 46: D303–D307.

Bohnsack KE, Hobartner C, Bohnsack MT. 2019. Eukaryotic 5-methylcytosine (m(5)C) RNA Methyltransferases: Mechanisms, Cellular Functions, and Links to Disease. Genes (Basel) 10.

Bolger AM, Lohse M, Usadel B. 2014. Trimmomatic: a flexible trimmer for Illumina sequence data. Bioinformatics 30: 2114–2120.

Burgess A, David R, Searle IR. 2016. Deciphering the epitranscriptome: A green perspective. J Integr Plant Biol 58: 822–835.

Casella G, Tsitsipatis D, Abdelmohsen K, Gorospe M. 2019. mRNA methylation in cell senescence. Wiley Interdiscip Rev RNA 10: e1547.

Chan PP, Lowe TM. 2016. GtRNAdb 2.0: an expanded database of transfer RNA genes identified in complete and draft genomes. Nucleic Acids Research 44: D184–D189.

Chen X, Li A, Sun BF, Yang Y, Han YN, Yuan X, Chen RX, Wei WS, Liu Y, Gao CC et al. 2019. 5-methylcytosine promotes pathogenesis of bladder cancer through stabilizing mRNAs. Nat Cell Biol 21: 978–990.

Cheng JX, Chen L, Li Y, Cloe A, Yue M, Wei J, Watanabe KA, Shammo JM, Anastasi J, Shen QJ et al. 2018. RNA cytosine methylation and methyltransferases mediate chromatin organization and 5-azacytidine response and resistance in leukaemia. Nat Commun 9: 1163.

Chi L, Delgado-Olguin P. 2013. Expression of NOL1/NOP2/sun domain (Nsun) RNA methyltransferase family genes in early mouse embryogenesis. Gene Expr Patterns 13: 319–327.

Clancy JL, Nousch M, Humphreys DT, Westman BJ, Beilharz TH, Preiss T. 2007. Methods to analyze microRNA-mediated control of mRNA translation. Methods Enzymol 431: 83–111.

Courtney DG, Chalem A, Bogerd HP, Law BA, Kennedy EM, Holley CL, Cullen BR. 2019a. Extensive Epitranscriptomic Methylation of A and C Residues on Murine Leukemia Virus Transcripts Enhances Viral Gene Expression. MBio 10.

Courtney DG, Tsai K, Bogerd HP, Kennedy EM, Law BA, Emery A, Swanstrom R, Holley CL, Cullen BR. 2019b. Epitranscriptomic Addition of m(5)C to HIV-1 Transcripts Regulates Viral Gene Expression. Cell Host Microbe 26: 217–227 e216.

Cui X, Liang Z, Shen L, Zhang Q, Bao S, Geng Y, Zhang B, Leo V, Vardy LA, Lu T et al. 2017. 5-Methylcytosine RNA Methylation in Arabidopsis Thaliana. Mol Plant 10: 1387–1399.

David R, Burgess A, Parker B, Li J, Pulsford K, Sibbritt T, Preiss T, Searle IR. 2017. Transcriptome-Wide Mapping of RNA 5-Methylcytosine in Arabidopsis mRNAs and Noncoding RNAs. Plant Cell 29: 445–460.

Delatte B, Wang F, Ngoc LV, Collignon E, Bonvin E, Deplus R, Calonne E, Hassabi B, Putmans P, Awe S et al. 2016. RNA biochemistry. Transcriptome-wide distribution and function of RNA hydroxymethylcytosine. Science 351: 282–285.

Desrosiers R, Friderici K, Rottman F. 1974. Identification of methylated nucleosides in messenger RNA from Novikoff hepatoma cells. Proc Natl Acad Sci U S A 71: 3971–3975.

Dominissini D, Moshitch-Moshkovitz S, Schwartz S, Salmon-Divon M, Ungar L, Osenberg S, Cesarkas K, Jacob-Hirsch J, Amariglio N, Kupiec M et al. 2012. Topology of the human and mouse m6A RNA methylomes revealed by m6A-seq. Nature 485: 201–206.

Du H, Zhao Y, He J, Zhang Y, Xi H, Liu M, Ma J, Wu L. 2016. YTHDF2 destabilizes m(6)A-containing RNA through direct recruitment of the CCR4-NOT deadenylase complex. Nat Commun 7: 12626.

Dubin DT, Taylor RH. 1975. The methylation state of poly A-containing messenger RNA from cultured hamster cells. Nucleic Acids Res 2: 1653–1668.

Edelheit S, Schwartz S, Mumbach MR, Wurtzel O, Sorek R. 2013. Transcriptome-wide mapping of 5-methylcytidine RNA modifications in bacteria, archaea, and yeast reveals m5C within archaeal mRNAs. PLoS Genet 9: e1003602.

Evers M, Shafik A, Schumann U, Preiss T. 2016. RNAModR: Functional analysis of mRNA modifications in R. bioRxiv doi:10.1101/080051: 080051.

Fray RG, Simpson GG. 2015. The Arabidopsis epitranscriptome. Curr Opin Plant Biol 27: 17–21.

Goddard JP, Maden BE. 1976. Reaction of HeLa cell methyl-labelled 28S ribosomal RNA with sodium bisulphite: a conformational probe for methylated sequences. Nucleic Acids Res 3: 431–440.

Goddard JP, Schulman LH. 1972. Conversion of exposed cytidine residues to uridine residues in Escherichia coli formylmethionine transfer ribonucleic acid. J Biol Chem 247: 3864–3867.

Goodchild J, Fellner P, Porter AG. 1975. The determination of secondary structure in the poly(C) tract of encephalomyocarditis virus RNA with sodium bisulphite. Nucleic Acids Res 2: 887–895.

Helm M, Alfonzo JD. 2014. Posttranscriptional RNA Modifications: playing metabolic games in a cell’s chemical Legoland. Chem Biol 21: 174–185.

Huang T, Chen W, Liu J, Gu N, Zhang R. 2019. Genome-wide identification of mRNA 5-methylcytosine in mammals. Nat Struct Mol Biol 26: 380–388.

Huang W, Lan MD, Qi CB, Zheng SJ, Wei SZ, Yuan BF, Feng YQ. 2016. Formation and determination of the oxidation products of 5-methylcytosine in RNA. Chem Sci 7: 5495–5502.

Hussain S, Aleksic J, Blanco S, Dietmann S, Frye M. 2013a. Characterizing 5-methylcytosine in the mammalian epitranscriptome. Genome Biol 14: 215.

Hussain S, Sajini AA, Blanco S, Dietmann S, Lombard P, Sugimoto Y, Paramor M, Gleeson JG, Odom DT, Ule J et al. 2013b. NSun2-mediated cytosine-5 methylation of vault noncoding RNA determines its processing into regulatory small RNAs. Cell Rep 4: 255–261.

Jacob R, Zander S, Gutschner T. 2017. The Dark Side of the Epitranscriptome: Chemical Modifications in Long Non-Coding RNAs. Int J Mol Sci 18.

Kennedy EM, Courtney DG, Tsai K, Cullen BR. 2017. Viral Epitranscriptomics. J Virol 91.

Khoddami V, Cairns BR. 2013. Identification of direct targets and modified bases of RNA cytosine methyltransferases. Nat Biotechnol 31: 458–464.

Krueger F, Andrews SR. 2011. Bismark: a flexible aligner and methylation caller for Bisulfite-Seq applications. Bioinformatics 27: 1571–1572.

Kumar L, M EF. 2007. Mfuzz: a software package for soft clustering of microarray data. Bioinformation 2: 5–7.

Legrand C, Tuorto F, Hartmann M, Liebers R, Jacob D, Helm M, Lyko F. 2017. Statistically robust methylation calling for whole-transcriptome bisulfite sequencing reveals distinct methylation patterns for mouse RNAs. Genome Res 27: 1589–1596.

Linder B, Jaffrey SR. 2019. Discovering and Mapping the Modified Nucleotides That Comprise the Epitranscriptome of mRNA. Cold Spring Harb Perspect Biol 11.

Liu JZ, Yue YN, Han DL, Wang X, Fu Y, Zhang L, Jia GF, Yu M, Lu ZK, Deng X et al. 2014. A METTL3-METTL14 complex mediates mammalian nuclear RNA N-6-adenosine methylation. Nature Chemical Biology 10: 93–95.

Liu N, Pan T. 2015. RNA epigenetics. Transl Res 165: 28–35.

Lorenz R, Bernhart SH, Honer Zu Siederdissen C, Tafer H, Flamm C, Stadler PF, Hofacker IL. 2011. ViennaRNA Package 2.0. Algorithms Mol Biol 6: 26.

Marbaniang CN, Vogel J. 2016. Emerging roles of RNA modifications in bacteria. Curr Opin Microbiol 30: 50–57.

Meyer KD, Jaffrey SR. 2017. Rethinking m(6)A Readers, Writers, and Erasers. Annu Rev Cell Dev Bi 33: 319–342.

Meyer KD, Saletore Y, Zumbo P, Elemento O, Mason CE, Jaffrey SR. 2012. Comprehensive analysis of mRNA methylation reveals enrichment in 3’ UTRs and near stop codons. Cell 149: 1635–1646.

Miao Z, Xin N, Wei B, Hua X, Zhang G, Leng C, Zhao C, Wu D, Li J, Ge W et al. 2016. 5-hydroxymethylcytosine is detected in RNA from mouse brain tissues. Brain Res 1642: 546–552.

Motorin Y, Helm M. 2019. Methods for RNA Modification Mapping Using Deep Sequencing: Established and New Emerging Technologies. Genes (Basel) 10.

Nachtergaele S, He C. 2018. Chemical Modifications in the Life of an mRNA Transcript. Annu Rev Genet 52: 349–372.

Park JE, Yi H, Kim Y, Chang H, Kim VN. 2016. Regulation of Poly(A) Tail and Translation during the Somatic Cell Cycle. Mol Cell 62: 462–471.

Peer E, Rechavi G, Dominissini D. 2017. Epitranscriptomics: regulation of mRNA metabolism through modifications. Curr Opin Chem Biol 41: 93–98.

Perry RP, Kelley DE. 1974. Existence of methylated messenger RNA in mouse L cells. Cell 1: 37–42.

Phizicky EM, Alfonzo JD. 2010. Do all modifications benefit all tRNAs? FEBS Lett 584: 265–271.

Quinlan AR, Hall IM. 2010. BEDTools: a flexible suite of utilities for comparing genomic features. Bioinformatics 26: 841–842.

Ramaswami G, Li JB. 2016. Identification of human RNA editing sites: A historical perspective. Methods 107: 42–47.

Rieder D, Amort T, Kugler E, Lusser A, Trajanoski Z. 2016. meRanTK: methylated RNA analysis ToolKit. Bioinformatics 32: 782–785.

Robinson MD, McCarthy DJ, Smyth GK. 2010. edgeR: a Bioconductor package for differential expression analysis of digital gene expression data. Bioinformatics 26: 139–140.

Roundtree IA, He C. 2016. RNA epigenetics--chemical messages for posttranscriptional gene regulation. Curr Opin Chem Biol 30: 46–51.

Saletore Y, Meyer K, Korlach J, Vilfan ID, Jaffrey S, Mason CE. 2012. The birth of the Epitranscriptome: deciphering the function of RNA modifications. Genome Biol 13: 175.

Schaefer M, Kapoor U, Jantsch MF. 2017. Understanding RNA modifications: the promises and technological bottlenecks of the ‘epitranscriptome’. Open Biol 7.

Schaefer M, Pollex T, Hanna K, Tuorto F, Meusburger M, Helm M, Lyko F. 2010. RNA methylation by Dnmt2 protects transfer RNAs against stress-induced cleavage. Genes Dev 24: 1590–1595.

Shafik A, Schumann U, Evers M, Sibbritt T, Preiss T. 2016. The emerging epitranscriptomics of long noncoding RNAs. Biochim Biophys Acta 1859: 59–70.

Sharma S, Yang J, Watzinger P, Kotter P, Entian KD. 2013. Yeast Nop2 and Rcm1 methylate C2870 and C2278 of the 25S rRNA, respectively. Nucleic Acids Res 41: 9062–9076.

Shen L, Liang Z, Wong CE, Yu H. 2019. Messenger RNA Modifications in Plants. Trends Plant Sci 24: 328–341.

Shi HL, Wei JB, He C. 2019. Where, When, and How: Context-Dependent Functions of RNA Methylation Writers, Readers, and Erasers. Molecular Cell 74: 640–650.

Sibbritt T, Patel HR, Preiss T. 2013. Mapping and significance of the mRNA methylome. Wiley Interdiscip Rev RNA 4: 397–422.

Sibbritt T, Shafik A, Clark SJ, Preiss T. 2016. Nucleotide-Level Profiling of m(5)C RNA Methylation. Methods Mol Biol 1358: 269–284.

Sloan KE, Warda AS, Sharma S, Entian KD, Lafontaine DLJ, Bohnsack MT. 2017. Tuning the ribosome: The influence of rRNA modification on eukaryotic ribosome biogenesis and function. Rna Biol 14: 1138–1152.

Sommer S, Lavi U, Darnell JE, Jr. 1978. The absolute frequency of labeled N-6-methyladenosine in HeLa cell messenger RNA decreases with label time. J Mol Biol 124: 487–499.

Squires JE, Patel HR, Nousch M, Sibbritt T, Humphreys DT, Parker BJ, Suter CM, Preiss T. 2012. Widespread occurrence of 5-methylcytosine in human coding and non-coding RNA. Nucleic Acids Res 40: 5023–5033.

Sun Z, Xue S, Xu H, Hu X, Chen S, Yang Z, Yang Y, Ouyang J, Cui H. 2019. Effects of NSUN2 deficiency on the mRNA 5-methylcytosine modification and gene expression profile in HEK293 cells. Epigenomics-Uk 11: 439–453.

Trixl L, Lusser A. 2019. The dynamic RNA modification 5-methylcytosine and its emerging role as an epitranscriptomic mark. Wiley Interdiscip Rev RNA 10: e1510.

Wagih O. 2017. ggseqlogo: a versatile R package for drawing sequence logos. Bioinformatics 33: 3645–3647.

Wang W. 2016. mRNA methylation by NSUN2 in cell proliferation. Wiley Interdiscip Rev RNA 7: 838–842.

Wang X, Lu ZK, Gomez A, Hon GC, Yue YN, Han DL, Fu Y, Parisien M, Dai Q, Jia GF et al. 2014. N-6-methyladenosine-dependent regulation of messenger RNA stability. Nature 505: 117-+.

Wei Z, Panneerdoss S, Timilsina S, Zhu J, Mohammad TA, Lu ZL, de Magalhaes JP, Chen Y, Rong R, Huang Y et al. 2018. Topological Characterization of Human and Mouse m(5)C Epitranscriptome Revealed by Bisulfite Sequencing. Int J Genomics 2018: 1351964.

Xiang JF, Yang Q, Liu CX, Wu M, Chen LL, Yang L. 2018. N(6)-Methyladenosines Modulate A-to-I RNA Editing. Mol Cell 69: 126–135 e126.

Xue S, Xu H, Sun Z, Shen H, Chen S, Ouyang J, Zhou Q, Hu X, Cui H. 2019. Depletion of TRDMT1 affects 5-methylcytosine modification of mRNA and inhibits HEK293 cell proliferation and migration. Biochem Biophys Res Commun 520: 60–66.

Yang L, Perrera V, Saplaoura E, Apelt F, Bahin M, Kramdi A, Olas J, Mueller-Roeber B, Sokolowska E, Zhang W et al. 2019a. m(5)C Methylation Guides Systemic Transport of Messenger RNA over Graft Junctions in Plants. Curr Biol 29: 2465–2476 e2465.

Yang X, Yang Y, Sun BF, Chen YS, Xu JW, Lai WY, Li A, Wang X, Bhattarai DP, Xiao W et al. 2017. 5-methylcytosine promotes mRNA export - NSUN2 as the methyltransferase and ALYREF as an m(5)C reader. Cell Res 27: 606–625.

Yang Y, Wang L, Han X, Yang WL, Zhang M, Ma HL, Sun BF, Li A, Xia J, Chen J et al. 2019b. RNA 5-Methylcytosine Facilitates the Maternal-to-Zygotic Transition by Preventing Maternal mRNA Decay. Mol Cell 75: 1188–1202 e1111.

Yu G, Wang LG, Han Y, He QY. 2012. clusterProfiler: an R package for comparing biological themes among gene clusters. OMICS 16: 284–287.

Zaccara S, Ries RJ, Jaffrey SR. 2019. Reading, writing and erasing mRNA methylation. Nat Rev Mol Cell Biol 20: 608–624.

Zhang HY, Xiong J, Qi BL, Feng YQ, Yuan BF. 2016. The existence of 5-hydroxymethylcytosine and 5-formylcytosine in both DNA and RNA in mammals. Chem Commun (Camb) 52: 737–740.

Zheng G, Dahl JA, Niu Y, Fedorcsak P, Huang CM, Li CJ, Vagbo CB, Shi Y, Wang WL, Song SH et al. 2013. ALKBH5 is a mammalian RNA demethylase that impacts RNA metabolism and mouse fertility. Mol Cell 49: 18–29.

